# MitoScope: a comprehensive mitochondrial genome analysis workflow for long-read sequencing data

**DOI:** 10.64898/2026.04.13.718259

**Authors:** Christina Zakarian, Joshua D Smith, Chee Hong Wong, Christian D Frazar, Erica Ryke, Sean R McGee, Matthew Richardson, Jeffrey M Weiss, Katherine M Munson, Kendra Hoekzema, Taralynn Mack, Youngjun Kwon, Jeffrey Ou, Shane Neph, Min-Hwan Sohn, Anna Minkina, James T Bennett, Andrew B Stergachis, Evan E Eichler, Chia-Lin Wei

## Abstract

The mitochondrial genome (mtDNA), rich in repeats and prone to nuclear mitochondrial DNA segments (NUMTs), is implicated in cancer, metabolic syndromes, and neurodegeneration, yet short-read sequencing yields incomplete catalogs, mapping artifacts, and false heteroplasmies. We introduce MitoScope, a scalable end-to-end long-read mitochondrial analysis workflow that integrates mtDNA-read selection, heteroplasmic variant detection, assembly, copy number estimation, read-level haplotyping, and NUMT profiling. Applied to benchmarking data from the Somatic Mosaicism Across Human Tissues (SMaHT) Network, we reveal high sensitivity and precision compared to existing methods and successfully uncover known mitochondrial biology relevant for variant interpretation, aging, and human disease.

## Background

Mitochondria play a central role in cellular energy production, metabolic homeostasis, and stress-responsive signaling [1,2]. Their dysfunction contributes to diverse human diseases ranging from neurodegeneration and cancer to cardiometabolic syndromes [3–7]. The human mitochondrial genome (mtDNA) is a 16.5 kilobase circular molecule, encoding 13 oxidative phosphorylation proteins, 22 tRNAs, and 2 rRNAs [8]. Unlike the nuclear genome, mtDNA exists at tens to thousands of copies per cell, where its high mutation rate and multi-copy organization generate extensive intra-cellular and intra-tissue variation (heteroplasmy) that is biologically important and technically challenging to resolve. In most tissues, wild-type and mutant mtDNA molecules coexist, and shifts in heteroplasmy levels can modulate oxidative phosphorylation, influence cellular fitness, and contribute to late-onset phenotypes even in clinically normal donors [3]. Somatic mtDNA mutations have been shown to accumulate in an age-dependent manner [5,9], yet their prevalence and spectrum across diverse healthy human tissues remain incompletely defined. Recent mtDNA studies using large-scale population databases (UK Biobank, All of Us, gnomAD) have largely been restricted to the use of short-read sequencing data from blood samples [9,10], lacking representation of a more complete spectrum of heteroplasmic variation on a tissue-dependent basis.

Beyond mitochondrial genomes, nuclear mitochondrial DNA segments (NUMTs) embedded in the nuclear genome can reshape local chromatin, perturb gene regulation, and confound interpretation of both nuclear and mitochondrial variants in rare disease and cancer genomics [11–13]. NUMT integrations have been linked to genomic instability, clonal evolution, and age-associated mitochondrial dysfunction [14], yet most current knowledge derives from germline or population-level catalogs [11], rather than tissue-resolved somatic events. Consequently, the extent, dynamics, and functional impact of NUMT mosaicism across human tissues remain largely undefined, limiting the ability to distinguish benign background integrations from potentially pathogenic events and to recognize NUMT-associated instability as a feature of somatic mosaicism [15,16].

Traditional short-read sequencing struggles to resolve the full spectrum of mtDNA variation in the presence of NUMTs as short fragments cannot distinguish reads arising from the mitochondrial versus the nuclear genome. NUMTs can range in size from <50 bp to the full length of mtDNA while sharing high sequence identity with mtDNA [11]. With short reads, this often leads to mapping artifacts, misassemblies, false heteroplasmies, and incomplete catalogs of both mtDNA variants and NUMTs. Despite short-read-based surveys of large cohorts having cataloged germline mtDNA polymorphisms and thousands of NUMTs [11], somatic mtDNA single-nucleotide variants (SNVs), indels (insertions/deletions), structural variants, and tissue-mosaic NUMT diversity are likely underestimated, particularly at low heteroplasmy frequencies relevant for somatic mosaicism.

Long-read sequencing technologies can capture full-length circular mtDNA molecules, span repetitive segments, and distinguish mtDNA from NUMTs using genome alignment and epigenetic signatures, due to higher 5mC levels in nuclear versus mitochondrial DNA. Most mitochondrial analysis tools are optimized for short-read data [17], and existing long-read tools, like Mitorsaw and Himito [18,19], are restricted in platform support, throughput, or scalability, limiting their use in cohort-scale, tissue-resolved analyses of somatic mtDNA and NUMTs. As a result, there is a critical need for a robust, scalable long-read workflow that can jointly resolve mtDNA copy number (CN), heteroplasmic SNVs and structural variants, and NUMTs across diverse tissues and donors. Here we present MitoScope, a workflow for long-read-specific mitochondrial analysis with stringent selection of mtDNA reads, high-fidelity heteroplasmic variant calling, circular mtDNA assembly, comprehensive quality control, and systematic NUMT profiling. Applied to benchmarking HapMap cell mixtures and tissues from the Somatic Mosaicism across Human Tissues (SMaHT) Network [20–22], MitoScope achieves high sensitivity and precision, recovers well-established biological patterns of mitochondria including tissue-specific mtDNA CN, low-frequency pathogenic heteroplasmies, and major-arc deletions enriched for microhomology, and identifies NUMTs in long-read data that are absent from existing short-read catalogs—exposing pervasive, tissue-resolved somatic mtDNA and NUMTs in healthy human tissues.

## Results

### MitoScope: A scalable long-read workflow for mtDNA selection, variant detection, assembly, and NUMT profiling

In developing MitoScope, we used a combined approach of custom scripts and existing state-of-the-art tools to build a fully comprehensive workflow for long-read specific mtDNA and NUMT analysis. Although individual mtDNA tools exist addressing specific tasks such as assembly and variant calling, a major limitation of most currently available workflows is the focus on only a single mtDNA analysis task [19,23,24]. MitoScope integrates five different key modules: 1) mtDNA selection, 2) variant calling and annotation, 3) assembly generation, 4) quality control, and 5) optional NUMT profiling using a single containerized workflow optimized to carefully address NUMT contamination (**Additional file 1: Fig. S1**). This eliminates the need to use an assortment of several tools for comprehensive mtDNA analysis from long-read data. Furthermore, MitoScope is implemented using Nextflow and fully containerized using Singularity, enabling execution of the integrated analysis modules with versioned software dependencies. A full list of software and container versions used by MitoScope as well as expected outputs from the workflow are provided **(Additional file 2: Table S1-S2).** Long-read sequencing data can be provided in the form of FASTQ, BAM, or CRAM files, either raw or aligned, providing flexibility for the user. Computational resources can be configured through the Nextflow config file, with default limits of 16 CPUs and 256 GB memory. A representative cohort of 100 PacBio samples, averaging 30x coverage, completed in approximately 90 minutes without NUMT profiling and 210 minutes with NUMT profiling. The full workflow with example input data and commands are available through the GitHub repository (https://github.com/czakarian/mitoscope) [25].

The mtDNA selection module is designed to remove contaminating NUMTs and ensure downstream tasks such as variant calling and assembly consist only of true mtDNA using several rounds of filtration. We begin by initiating an alignment-independent, k-mer-based extraction of candidate mtDNA reads from either raw sequencing reads or chrM-mapped reads from aligned input. A k-mer count database of mtDNA k-mer sequences (k= 29) is used to selectively retain reads with >2,500 matching k-mers, prioritizing reads with sufficient length and sequence similarity to mtDNA. Reads selected by k-mer matching are then aligned to the mitochondrial genome using platform-specific Minimap2 presets (--map-ont for ONT and --map-hifi for PacBio), followed by filtration based on alignment and methylation signatures. mtDNA is known to exhibit very low levels of 5mC methylation relative to the nuclear genome [26], providing a useful secondary method for differentiation in addition to alignment. Reads that do not fully align to the mitochondrial genome or display elevated 5mC methylation are discarded (Methods).

We next built out a variant detection module with integrated gene and disease annotations that was inclusive to both small variants and larger structural variants and allowed for both read-level and bulk variant analysis. We found most existing mtDNA variant callers did not include structural variants and were solely focused on SNVs and indels, despite the known prevalence of large-scale mtDNA deletions in various disorders [3,7] and associations with age [27,28]. To address this limitation, we build a variant detection module encompassing SNVs, indels, and structural variants by utilizing baldur [29], a long-read mitochondrial variant caller for heteroplasmic SNVs, indels, and large deletions, alongside the long-read structural variant (SV) caller, Sniffles2 [30], with parameters customized to identify heteroplasmic SVs (Methods). For read-level variant analysis, a matrix is generated per sample indicating variant presence or absence across individual reads. Identified SNV and indel variants are annotated using Variant Effect Predictor (VEP) [31] and MITOMAP [32] for associated genes, functional consequence, pathogenicity, and population frequencies. Furthermore, for exploration of epigenetic patterns, methylation calls are generated from base modification signals (5mC, 5hmC) and processed using either pb-CpG-tools [33] for PacBio data and Modkit [34] for ONT data.

*De novo* circular mitochondrial genome assemblies are produced using meta-flye [35], a long-read assembler optimized for metagenomic contexts. To reduce computational burden and limit misassemblies associated with excessive coverage, each assembly attempt uses a random subset of 100 downsampled mtDNA reads. If an approximately 16.5 Kbp circular assembly is not obtained, meta-flye is rerun using a newly selected random subset of 100 reads. This process is repeated until a valid circular assembly is generated or the maximum number of attempts is reached (default = 10). At the same time, quality metrics are generated per sample including read counts from each stage of mtDNA selection, length distributions (histograms/median), mtDNA genome coverage calculated using Mosdepth [36], mtDNA CN, and mitochondrial haplogroups assigned by Haplogrep3 [37].

To define the extent and dynamics of NUMT integration across human tissues, MitoScope offers an optional module to perform systematic characterization of non-reference NUMTs using long-read data. Nuclear insertions that align to the mitochondrial genome or split reads with supplementary alignments to both the mitochondrial and nuclear genomes are reported as NUMTs with their corresponding nuclear and mitochondrial breakpoint locations, size, and levels of read support (Methods).

### MitoScope achieves accurate, high-fidelity variant calling in HapMap mixture benchmarking samples

The quantitative performance of MitoScope in variant calling was evaluated using genetically diverse cell mixtures constructed from six selected lymphoblast cell lines of the International HapMap project mixed at defined ratios (83.5% HG005, 10% HG02622, 2% HG02486, 2% HG02257, 2% HG002, 0.5% HG00438) (**Fig. 1A**). This publicly available benchmarking resource was specially designed and carried out through the efforts of the SMaHT Network [20–22] to generate variant allele fractions from 0.25% to 16.5%, approximating somatic mosaicism in tissues and serving as analytical ground truth for sensitivity, precision, and limit of detection across variant classes. Using MitoScope, we processed sequencing data (n=7) from Pacific Biosciences (PacBio) and Oxford Nanopore Technologies (ONT) generated across five SMaHT genome characterization centers (GCCs: Baylor College of Medicine (BCM), University of Washington and Seattle Children Research Institute (UW-SCRI), Washington University at St. Louis and Van Andell Institute (WashU-VAI), Broad Institute (Broad), and New York Genome Center (NYGC)) and determined coverage, median read length, and copy number (**Additional file 2: Table S3**). Matched short-read data (n=4) generated across SMaHT GCCs (**Additional file 2: Table S4**) was incorporated to investigate long-read and short-read platform-specific differences in mitochondrial copy number and variant calling results.

**Fig. 1.**
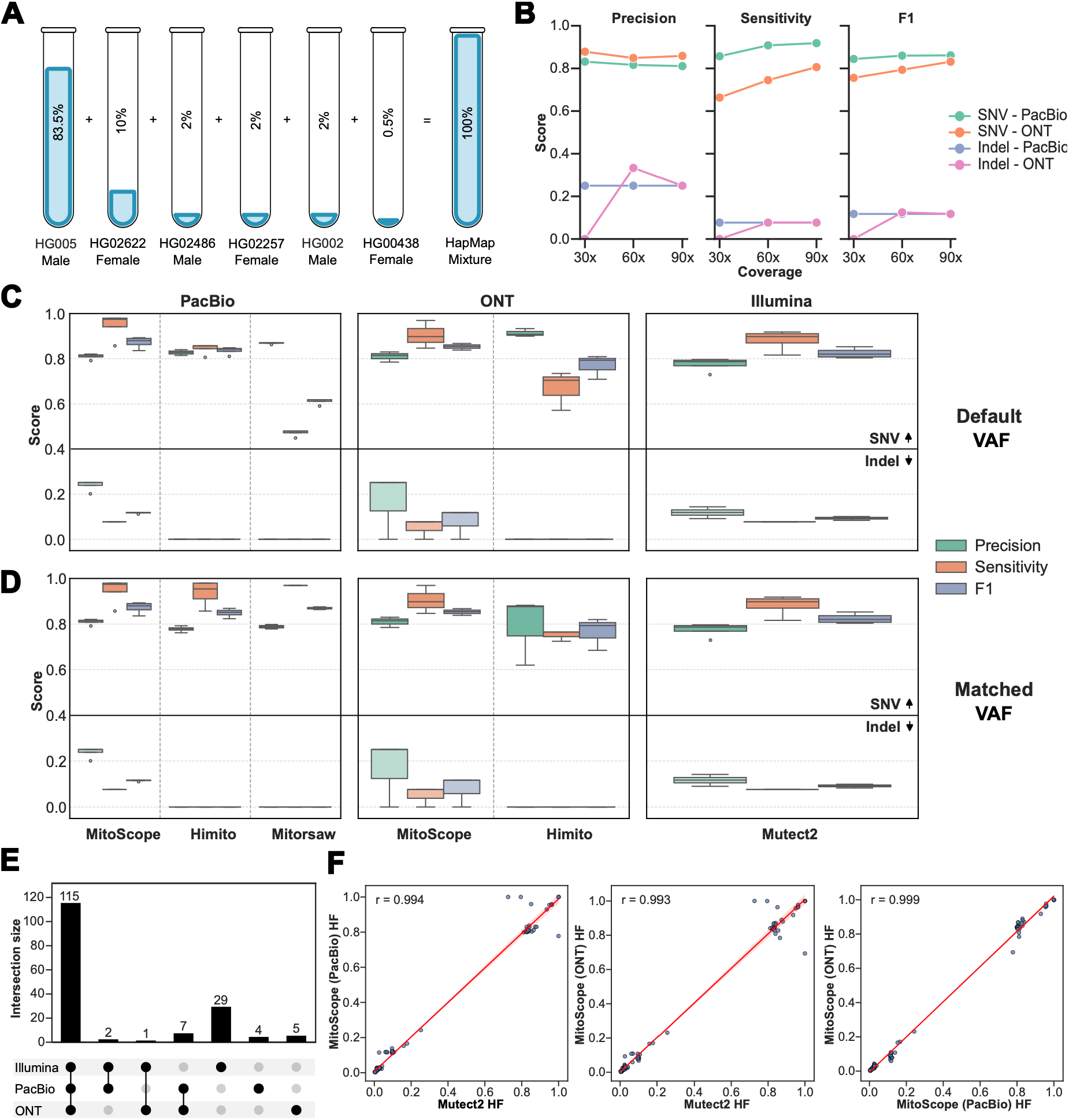
MitoScope achieves accurate, high-fidelity variant calling in HapMap mixture benchmarking samples. **A** Tool performance was evaluated using the HapMap mixture cell line data generated by the SMaHT network. The HapMap mixture was made using six samples from the International HapMap project at varying mixture ratios. **B** SNV and indel variant calling performance across varying coverage levels (30x, 60x, 90x) using downsampled PacBio and ONT HapMap samples. **C-D** Precision, sensitivity, and F1 scores were calculated for SNV and indel variant calling results from MitoScope and other long-read (Himito, Mitorsaw) and short-read (Mutect2) mitochondrial variant callers using a truth set (98 SNVs, 13 indels) derived from HPRC pangenome assemblies. Tools were run using their default settings (**C**) and separately using a matched minimum VAF threshold of 0.0025 (**D**). Data are represented as box- and-whisker with the box representing interquartile range (IQR, 25th–75th percentiles), the line within the box representing the median, whiskers extending to 1.5*IQR, and outliers depicted using individual points (n=3 GCC replicates for ONT, n=4 for PacBio, and n=4 for Illumina). **E** Concordance of HapMap variants across MitoScope PacBio and ONT and Mutect2 Illumina calls. **F** Concordance was measured for heteroplasmy frequencies (HF) of 115 variants reported by both MitoScope (PacBio, ONT) and Mutect2 (Illumina). A Pearson correlation coefficient (r) was calculated for each pair of comparisons.

Notably, mtDNA abundance inferred from the long-read data was highly variable, ranging from 18 to 157 copies per diploid nuclear genome and considerably lower than the mtDNA CN inferred from short-read data (CN: 328-1071). mtDNA CN inferred from long-read datasets displayed an overall reverse correlation with median read length (**Additional file 1: Fig. S2**) and was associated with variation in the library preparation protocols deployed by different GCCs. Long-read sequencing data generated by protocols optimized for longer read lengths (median read length >=20 Kbp) with size selection of the sheared DNA severely underestimated the mtDNA CN, consistent with prior reports of significant associations of library fragmentation and size selection approaches on relative mtDNA abundance [38]. In standard long-read datasets, mtDNA abundance can therefore be underestimated, as mtDNA is preferentially lost during DNA shearing and long fragment selection. Furthermore, because PacBio Fiber-seq [39] deploys nuclei preparation which depletes mitochondria not physically attached to the nuclear membrane, protocols using Fiber-seq will have reduced recoverable mtDNA in long-read libraries.

To evaluate variant detection, MitoScope SNV and indel calls were evaluated against a mitochondrial variant truth set derived from the Human Pangenome Reference Consortium (HPRC) version 2 pangenome-graph variant calls from the minigraph-cactus pipeline [40,41]. We subset the HPRC calls to chrM variants present in at least one of the six HapMap cell lines (HG005, HG02622, HG02486, HG02257, HG002, and HG00438). Because the HapMap mixture sample is a pooled mixture and individual sample origins cannot be distinguished in the sequencing data, genotype information from the six component cell lines was collapsed to create a single presence/absence truth set containing 98 SNVs and 13 indels.

We examined the effect of the minimum variant allele frequency (VAF) threshold on MitoScope variant-calling performance on PacBio and ONT data using the truth set (**Additional File 1: Fig. S3**). For SNVs, we tested minimum VAF thresholds of 1%, 0.5%, 0.25%, and 0.1%. For indels, we evaluated thresholds of 10%, 5%, and 1% as long-read sequencing is more prone to indel artifacts, particularly in homopolymer and low-complexity regions [42,43]. For SNVs, lowering the threshold from 0.5% to 0.25% increased sensitivity while maintaining precision, whereas reduction to 0.1% provided minimal additional sensitivity. For indels, sensitivity and precision were overall much lower than for SNVs. The 10% threshold yielded the highest precision, with little sensitivity gain at 5%. Lowering the threshold to 1% provided a modest increase in sensitivity but substantially reduced precision. With performance between PacBio and ONT being largely comparable, we used minimum VAF thresholds of 0.25% for SNVs and 10% for indels in all subsequent analyses, prioritizing sensitivity for low heteroplasmy SNVs while limiting false-positive indels.

Using the established VAF thresholds, we next explored MitoScope performance on PacBio and ONT data from the BCC GCC across varying coverage levels for SNVs and indels **(Fig. 1B).** Sensitivity and precision from 30x, 60x, and 90x coverage data were evaluated. For both sequencing platforms, sensitivity for SNVs gradually increased with increasing coverage, whereas precision remained relatively stable. PacBio data showed moderately higher sensitivity compared to ONT data at equal coverage, while precision between platforms was more comparable and slightly lower for PacBio data. Indel performance was overall much lower compared to SNVs as we observed earlier with a max observed F1-score of 0.125 compared to 0.86 for SNVs.

Variant calling performance of MitoScope was next evaluated alongside Himito and Mitorsaw, two long-read-specific mitochondrial variant calling tools, as well as high coverage short-read data processed by GATK’s Mutect2 [44] in mitochondrial mode (**Fig. 1C; Additional file 2: Table S5**). MitoScope was run using our established minimum VAF thresholds and remaining tools were initially run using their default settings. MitoScope’s performance in detecting SNVs from long reads was comparable to Himito and Mitorsaw in terms of precision with slightly higher levels of sensitivity, potentially owing to the lower minimum VAF threshold employed by MitoScope. Compared to Mutect2’s performance on high coverage short reads, MitoScope showed equally high sensitivity and precision for SNV detection across both PacBio and ONT data. Indel detection showed low sensitivity and precision across all callers and sequencing technologies, suggesting that the reduced indel performance was not specific to MitoScope, but more indicative of discordance in indels called directly from long- and short-read whole genome sequencing (WGS) data and those identified from the pangenome assemblies. To investigate the basis of the discrepancy, we further examined indels present in the pangenome-derived variant calls that were not detected in either long- or short-read WGS data. Of the 11 discrepant indels, 9 originated from a single HapMap sample, HG02486. Notably, 8 of these 9 indels were concentrated within a 115-bp region (chrM:14710-14825) and were relatively larger in size (>8bp). The clustering and size of these indels combined with their absence in both short and long read data could suggest an assembly-related artifact in the HG02486 mitochondrial assembly rather than true biological variation.

To account for potential differences in sensitivity and precision due to differences in the default minimum VAF thresholds enforced by different callers, we also compared performance using matched VAF thresholds (**Fig. 1D; Additional file 2: Table S6**). Himito and Mitorsaw’s minimum VAF thresholds were lowered from 0.01 and 0.10, respectively, to match MitoScope’s lower threshold of 0.0025. Mutect2 parameters were left unchanged since the tool does not have an adjustable minimum VAF setting. For SNVs, performance across tools on PacBio data was nearly identical whereas MitoScope continued to show higher sensitivity compared to Himito on ONT data. Performance on indels did not show improvement across any tools. Sample-to-sample variation between GCC replicates may reflect a combination of library preparation, coverage, and other sample-processing differences.

Platform-specific concordance of specific variant calls and reported heteroplasmy levels was next assessed across PacBio, ONT, and Illumina data (**Fig. 1E and F**). Most variants were shared across all three platforms (n=115) including variants with heteroplasmy levels as low as 0.3%. An additional 29, 5, and 4 variants were unique to Illumina, ONT, and PacBio, respectively. These platform-specific calls were enriched for low frequency heteroplasmic variants, indels, and variants in homopolymer regions. Among the Illumina-specific calls, 13 were indels, with heteroplasmy levels ranging from 0.1%-77%. In comparison, ONT identified no unique indels, while PacBio identified one unique indel at 10.7%, just above the minimum indel VAF threshold employed by MitoScope. Heteroplasmy levels of long-read MitoScope variant calls were highly concordant with the short-read Mutect2 variant calls (**Fig. 1F**; **Additional file 2: Table S7**). Using variants detected across all three platforms, we compared mean heteroplasmy levels across GCCs for each shared variant. Heteroplasmy levels were highly concordant between the PacBio and ONT calls made by MitoScope (Pearson correlation coefficient R>0.99) and between the Mutect2 short-read calls against both the MitoScope PacBio and ONT calls (R>0.99). A discordant call was identified in a homopolymer region (310T>TC) where Mutect2 called the indel homoplasmic and both PacBio and ONT reported heteroplasmy levels between 70-80%. Two other discordant calls were identified (8860A>G, 4769A>G) where MitoScope called variants homoplasmic on both PacBio and ONT data, while Mutect2 called the same variants heteroplasmic with frequencies between 70-80%. Manual inspection of short and long reads using pileup data supported the homoplasmic calls for the discordant variants (**Additional file 1: Fig. S4**). Despite variant presence in the population at high frequencies (>97% in MITOMAP) and across haplogroups, both variants have previously been reported as being miscalled by Mutect2 [45], corroborating the importance of the high accuracy calls from MitoScope. Together, these results indicate that while most mitochondrial variants are highly concordant across short and long-read platforms, discordant calls are more frequently associated with low frequency heteroplasmies, indels, and calls in homopolymer regions, features that are more consistent with potential sequencing artifacts.

### Platform-specific differences in mtDNA copy number and variant detection

As a pilot application of MitoScope, we next profiled mitochondrial genome variation across four phenotypically normal postmortem donors (ST001-ST004) collected as part of benchmarking efforts in the SMaHT Network, examining somatic mosaicism in four tissues (i.e., lung, liver, colon, and brain) (**Fig. 2A**). Donors were male, spanning young (<25 years; ST001 and ST003) and aged (>70 years; ST002 and ST004) age groups. We processed PacBio and ONT sequencing data generated at each of the five GCCs using MitoScope (**Additional file 2: Table S8**) alongside available matched bulk Illumina WGS data to compare long-read and short-read performance on mtDNA.

**Fig. 2.**
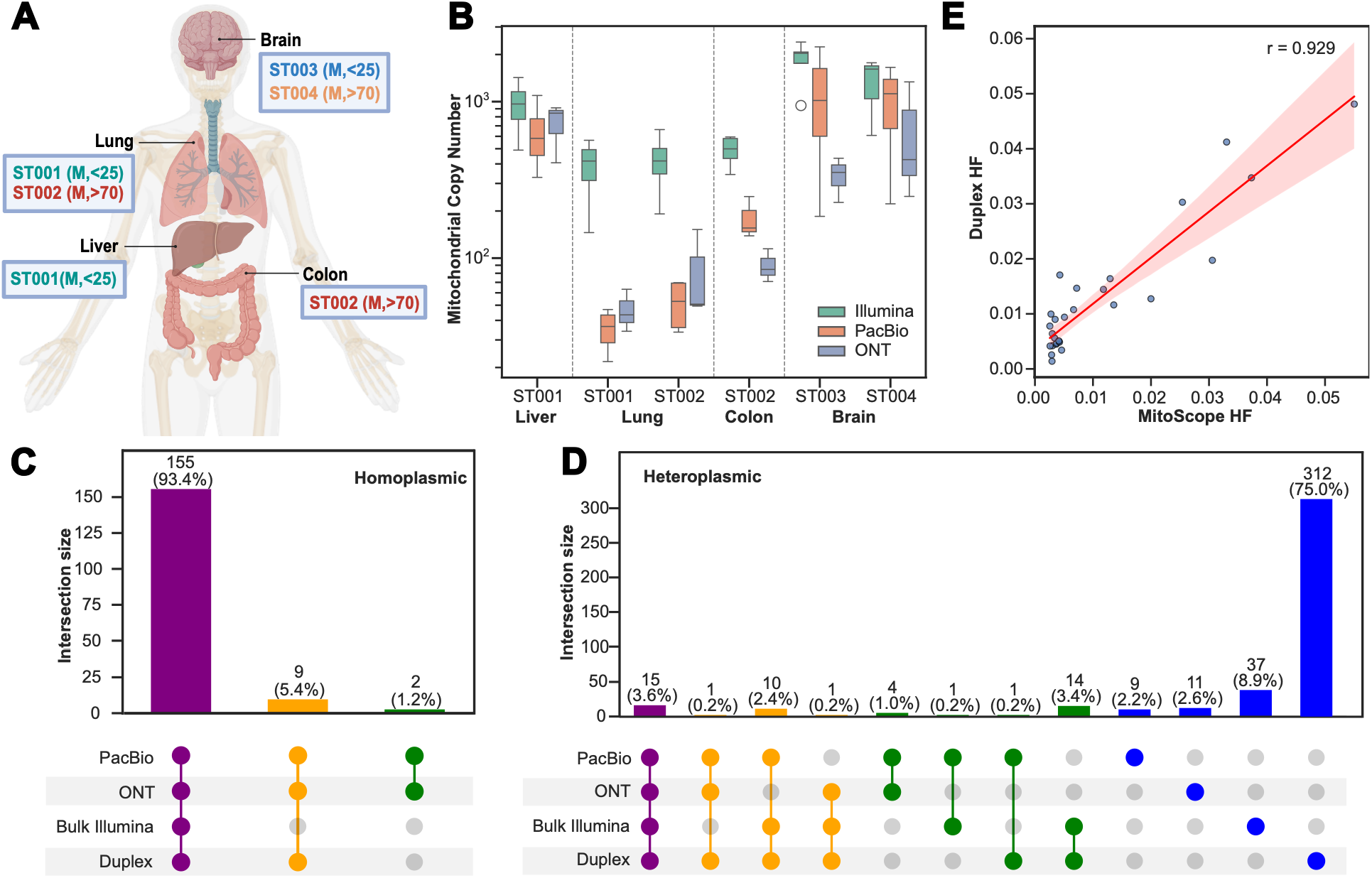
Platform-specific differences in mtDNA copy number and variant detection. **A** Overview of the SMaHT benchmarking tissues and donors with sex and age indicated in parenthesis. **B** Mitochondrial copy number estimates from PacBio, ONT, and Illumina sequencing data. Data are represented as box-and-whisker with the box representing interquartile range (IQR, 25th–75th percentiles), the line within the box representing the median, and whiskers extending to 1.5*IQR (n=3 GCC replicates for each ONT sample; n=4 GCC replicates each for PacBio ST001 Liver, ST001 Lung, ST002 Lung; and n=3 for PacBio ST002 Colon, ST003 Brain, ST004 Brain; n=4 GCC replicates for ST001 Liver and n=5 GCC replicates for remaining samples). **C-D** Upset plots for (**C**) homoplasmic and (**D**) heteroplasmic variant calls showing the distribution of concordant and discordant calls made across PacBio and ONT (using MitoScope), bulk short-read Illumina (using Mutect2), and duplex sequencing data (processed using CODECsuite and pileup variant calls). **E** Concordance of heteroplasmy frequencies (HF) of the variants shared between MitoScope (averaged ONT and PacBio heteroplasmy) and the duplex data (n=29).

mtDNA copy-number estimates derived from PacBio and ONT data showed higher values in brain and liver than in lung and colon. Matched bulk Illumina WGS data showed a similar tissue ordering, although the fold differences were substantially smaller than those observed in long-read datasets (**Fig. 2B**). Our findings from long-read data corroborate the known mitochondrial enrichment in higher-energy demanding tissues such as brain, liver, and muscle [46]. Together with the HapMap-mixture analysis, these results indicate that long-read library preparation can underestimate mtDNA representation and amplify apparent tissue differences, particularly in samples with low mtDNA abundance. Therefore, long-read-derived copy-number estimates should be interpreted cautiously as protocol-sensitive relative measures rather than absolute estimates of tissue-specific mtDNA copy number.

To explore mitochondrial SNV and indel variation across donors and tissues, we evaluated variants detected by PacBio and ONT, considering SNVs and indels and homoplasmic and heteroplasmic variants separately. Across 245 total variant calls, 78% (n=192) were shared between long-read platforms with 32 unique to PacBio and 21 unique to ONT. Variants in homopolymer regions (chrM:302-311,16179-16183), more prone to sequencing artifacts, were largely platform-specific (6 shared, 10 PacBio-specific, 9 ONT-specific). Of the remaining variants outside of homopolymer regions, nearly all were SNVs (166 homoplasmic, 53 heteroplasmic) with one heteroplasmic indel that was only called in PacBio data and at a heteroplasmy level just above MitoScope’s minimum VAF threshold for indels (>10%). All homoplasmic and 38% (n=20) of heteroplasmic SNVs were shared between PacBio and ONT. Considering the lack of indel calls and overall higher discordance of variants observed within homopolymer regions, we focused our subsequent analyses on SNVs outside of homopolymer regions.

To comprehensively evaluate SNVs detected by MitoScope, we next performed a multi-platform analysis using matched bulk short-read and duplex sequencing data. We used matched duplex sequencing data generated by the SMaHT benchmarking effort as an orthogonal platform with per-base error rates substantially lower than can be obtained with conventional WGS [47]. We found this approach especially valuable for evaluating lower frequency heteroplasmic variants (<1%) identified in the long-read data, which can be difficult to distinguish from background sequencing errors. Duplex-sequencing generates consensus read sequences by independently sequencing both strands (Watson and Crick) of the same DNA molecule and requiring concordance between the strands, reducing error rates and increasing sensitivity [47]. Despite its extremely high sensitivity, duplex sequencing is often limited by uneven and overall lower coverage compared to bulk WGS [47]. To overcome this potential limitation, we used the matched, high coverage (1289-1789x) bulk short-read Illumina data as additional support for low-frequency heteroplasmic variants.

Surprisingly, of the 166 homoplasmic SNVs common to PacBio and ONT, two were not called in duplex and bulk short-read data, with another nine missing from the bulk short-read data (**Fig. 2C**). For the missing duplex calls, we identified coverage dropout with no more than 20 total reads covering the relevant sites. Examination of the nine other missing homoplasmic calls in bulk short-read data revealed the same variant that was miscalled by Mutect2 in the short-read HapMap sample (8860A>G) and several other miscalled variants (225G>A, 226T>C, 5147G>A, 7424A>G), suggesting a caller-specific issue. Across heteroplasmic variants, we observed much more discordance between platforms. Duplex revealed the greatest number of unique calls (n=312) followed by bulk short-read data (n=37) and far less in ONT (n=11) and PacBio (n=9) (**Fig. 2D**). We identified 15 heteroplasmic variants that were shared across all 4 platforms, 12 shared across three platforms, and another 20 shared across two platforms. Of the heteroplasmic SNVs detected by MitoScope, 55% (n=29) were supported by duplex and/or bulk short-read data, with concordant heteroplasmy estimates spanning 0.25% to 5% (**Fig. 2E**). The remaining platform-specific long-read calls, 11 ONT-specific and 9 PacBio-specific heteroplasmic SNVs, were not supported by either short-read assay and were therefore classified as lower confidence. Their lack of confirmation may reflect platform-specific noise, stochastic sampling, or insufficient site-specific coverage in the orthogonal datasets. Interestingly, the large number of heteroplasmic variants unique to duplex data were all at low heteroplasmy levels (<1.5%) (**Additional File 1: Fig. S5**), revealing a potentially large reservoir of somatic mitochondrial variants undiscoverable by bulk long-read and short-read WGS.

### MitoScope uncovers established biological patterns of mtDNA variation across human tissues

For further analysis of tissue and donor-specific variation, we retained long-read variant calls from MitoScope that were called by at least two different sources (PacBio, ONT, bulk short-reads, duplex). Lower confidence variants called by MitoScope that were observed only once in either PacBio or ONT data were omitted from further downstream analysis. This filtration strategy resulted in a total of 166 homoplasmic and 33 heteroplasmic high-confidence SNVs, enabling robust inclusion and characterization of low-frequency heteroplasmies (HF<1%) that would otherwise be difficult to distinguish from technical noise when reliant purely on long-read data.

High-confidence SNVs across donors and tissues revealed a distinct bimodal distribution, consistent with prior observed patterns of mtDNA mutational spectra [9], predominantly homoplasmic, with a second peak of heteroplasmic variants, of which 24 were ultra-low frequency (<1%), 8 at 1–5%, and 1 at 5.7% (**Fig. 3A; Additional file 2: Table S9**). Of the 27 unique heteroplasmic SNVs detected in our limited survey, a missense variant (14889G>A) in *MT-CYB* present at a heteroplasmy level of 0.38%, has not been captured in MITOMAP, a human mitochondrial genome database with an extensive collection of mitochondrial SNVs (n=19,973), reinforcing the value of somatic tissue-based analysis to uncover somatic mosaicism in mtDNA.

**Fig. 3.**
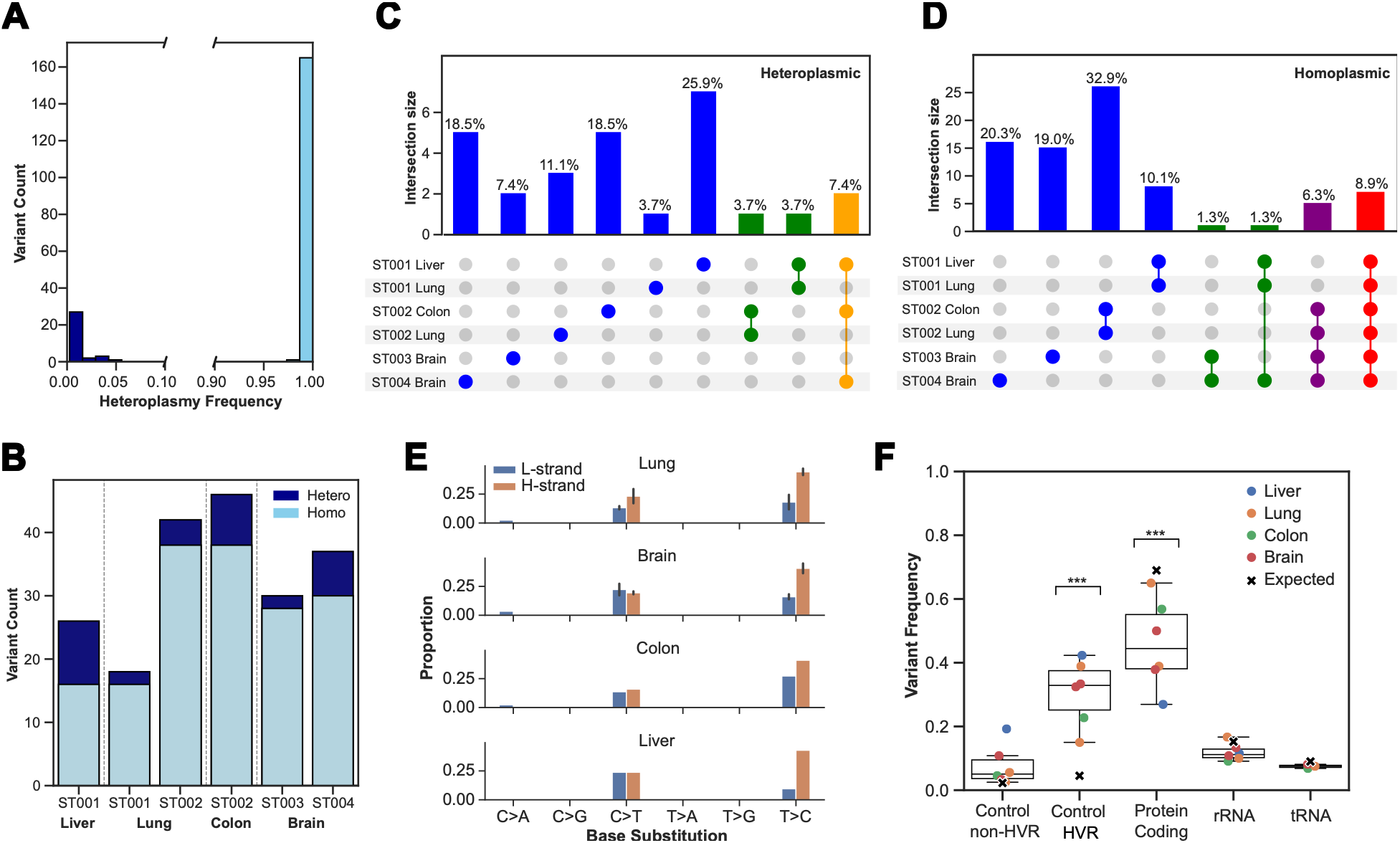
MitoScope uncovers established biological patterns of mtDNA variation across human tissues. **A** Distribution of heteroplasmy frequencies for high-confidence SNVs across all combined tissues and donors. **B** Heteroplasmic and homoplasmic SNV counts per tissue and donor. **C-D** Upset plots for (**C**) heteroplasmic and (**D**) homoplasmic variants showing the distribution of unique and common variants across donors and tissues. **E** SNVs categorized by base substitution on light and heavy strands. **F** SNV frequencies across the non-hypervariable (non-HVR) and hypervariable (HVR) control region, protein coding genes, rRNA genes, and tRNA genes were normalized by length and compared with length-based expectations (***P < 0.001, FDR-adjusted binomial test).

Homoplasmic SNVs were shared among tissues from the same donor, indicating their germline nature while the numbers and identities of heteroplasmic SNVs varied substantially across donors and tissues, suggesting they were mostly somatic (**Fig. 3B**). ST002 Lung and ST004 Brain had more heteroplasmic variants than matched tissues from younger donors, ST001 Lung and ST003 Brain (**Fig. 3B**), consistent with the well-established age-associated accumulation of mtDNA mutations [9,48,49]. Base substitutions were predominantly transitions and showed consistent H-strand bias across tissues (**Fig. 3E**), a pattern that has been linked to replication-linked errors and the increased susceptibility of the H-strand to deamination during asymmetric replication [9]. Across the mitochondrial genome, hypervariable segments of the control region showed significant SNV enrichment (∼6.3-fold; P < 0.001), whereas protein-coding regions showed significant depletion (∼1.4-fold; P < 0.001) **(Fig 3F)**. SNV frequencies in the non-hypervariable control region and in rRNA and tRNA genes were closer to length-based expectations. This distribution recapitulates established regional patterns of mtDNA variation, with greater polymorphism in hypervariable control-region sites and greater constraint in protein-coding and RNA genes [50], demonstrating MitoScope’s ability to recover known features of mitochondrial genome variation.

Using both homoplasmic and heteroplasmic SNVs, we next quantified unique and shared variants across donors and tissues. Homoplasmic variants, presumed germline origin, were predominantly donor-specific (82%), with only a small subset shared between donors, consistent with a limited pool of common mtDNA polymorphisms **(Fig. 3C)**. In contrast, 85% of heteroplasmic variants were unique in individual tissues and donors, indicating extensive, individualized somatic mosaicism across the mitochondrial genome **(Fig. 3D)** with rare heteroplasmic variants shared between tissues of the same donor likely representing either maternally inherited heteroplasmies or early somatic events that arose before tissue divergence. Heteroplasmic variants between donors with the same tissue type (ST001 and ST002 Lung, ST003 and ST004 Brain) were not observed.

Across tissues, multiple mtDNA SNVs mapped to MITOMAP entries with reported or confirmed disease associations (**Additional File 1: Fig. S6**). MITOMAP classifies variants as “reported,” with “conflicting reports,” or as “confirmed” (Cfrm) disease variants, where the latter is further annotated according to the ClinGen Pathogenicity Ratings (VUS - Variant of Uncertain Significance; LP - Likely Pathogenic; P - Pathogenic), reflecting the cumulative strength and consistency of clinical and functional evidence. Notably, each tissue harbored several reported variants (published in literature with limited evidence). Among these, we detected the canonical pathogenic m.3243A>G substitution in *MT-TL1* (MELAS-associated) at 2.5% heteroplasmy in ST004 Brain and a pathogenic m.10197G>A missense variant in *MT-ND3* linked to Leigh Disease at 0.35% heteroplasmy in ST002 Lung. These low-frequency pathogenic variants were further validated in the duplex data at 3% and 0.9% heteroplasmy levels, respectively. Using MitoScope, we show that we can successfully uncover low-frequency, disease-associated heteroplasmies in clinically unremarkable postmortem tissues, consistent with previous reports that pathogenic and disease-associated mtDNA variants can occur at low levels in otherwise healthy individuals [51,52], reflecting MitoScope’s ability to detect established patterns of somatic mtDNA variation.

### MitoScope enables read-level profiling of mitochondrial variant co-occurrence

MitoScope generates per-sample read-by-variant matrices that record the presence or absence of SNVs and indels on individual mtDNA molecules, together with a separate read-level file containing large deletions greater than 45 bp. These outputs enable direct assessment of molecular haplotypes and co-occurrence between SNVs, indel and large deletions. To illustrate this capability, we analyzed heteroplasmic variants in the ST004 Brain (**Additional File 1: Fig. S7**). We selected five heteroplasmic variants (VAF ranging from 0.5% to 5%) detected by both PacBio and ONT technologies and retained reads spanning all five variant positions. Across 71,643 combined PacBio and ONT reads, 87.6% carried only reference alleles at the evaluated sites. Among the 8417 reads carrying >=1 heteroplasmic variant, 96.6% (8,136 reads) carried one variant, 3.3% (278 reads) carried two variants, and 0.04% (3 reads) carried three variants. In addition, 52 of 8,417 variant-containing reads (0.6%) contain at least one heteroplasmic SNV and a large deletion (>45bp). Therefore, most variant-containing reads carried a single heteroplasmic variant, whereas a small fraction of reads showed co-occurrence of multiple variants on the same read. Although the low heteroplasmy levels and limited read counts prevent inferring clonal relationships, these data demonstrate that MitoScope supports direct read-level characterization of mtDNA molecular haplotypes and co-occurrence patterns.

### MitoScope uncovers age- and tissue-associated patterns of mitochondrial deletion landscapes

Large-scale deletions in the mitochondrial genome have been extensively characterized in the context of aging research and more recently implicated in primary mitochondrial disease, neurodegenerative disorders, and multisystem syndromes [3,7,27,28]. Here, we used MitoScope structural variant calls to systematically map the deletion landscape across benchmarking tissues. We observed widespread deletions spanning hundreds to thousands of base pairs, with the vast majority localizing to the major arc, which sits between the origins of replication of the light and heavy strand, and contains most protein-coding genes (**Fig. 4A**). Despite platform- and center-specific variation in absolute deletion frequencies, deletions were consistently most abundant in brain, with the aged brain (ST004) showing >2-fold higher total deletion heteroplasmy than the younger brain (ST003; HF 1.22% vs 0.45%), and a similar trend in aged versus young lung (ST002 vs ST001; HF 0.26% vs 0.19%) (**Fig. 4B**). Together, these findings are consistent with the recognized age-associated accumulation of somatic mtDNA deletions in the human brain, a phenomenon that has been extensively documented in aging research [28,53].

**Fig. 4.**
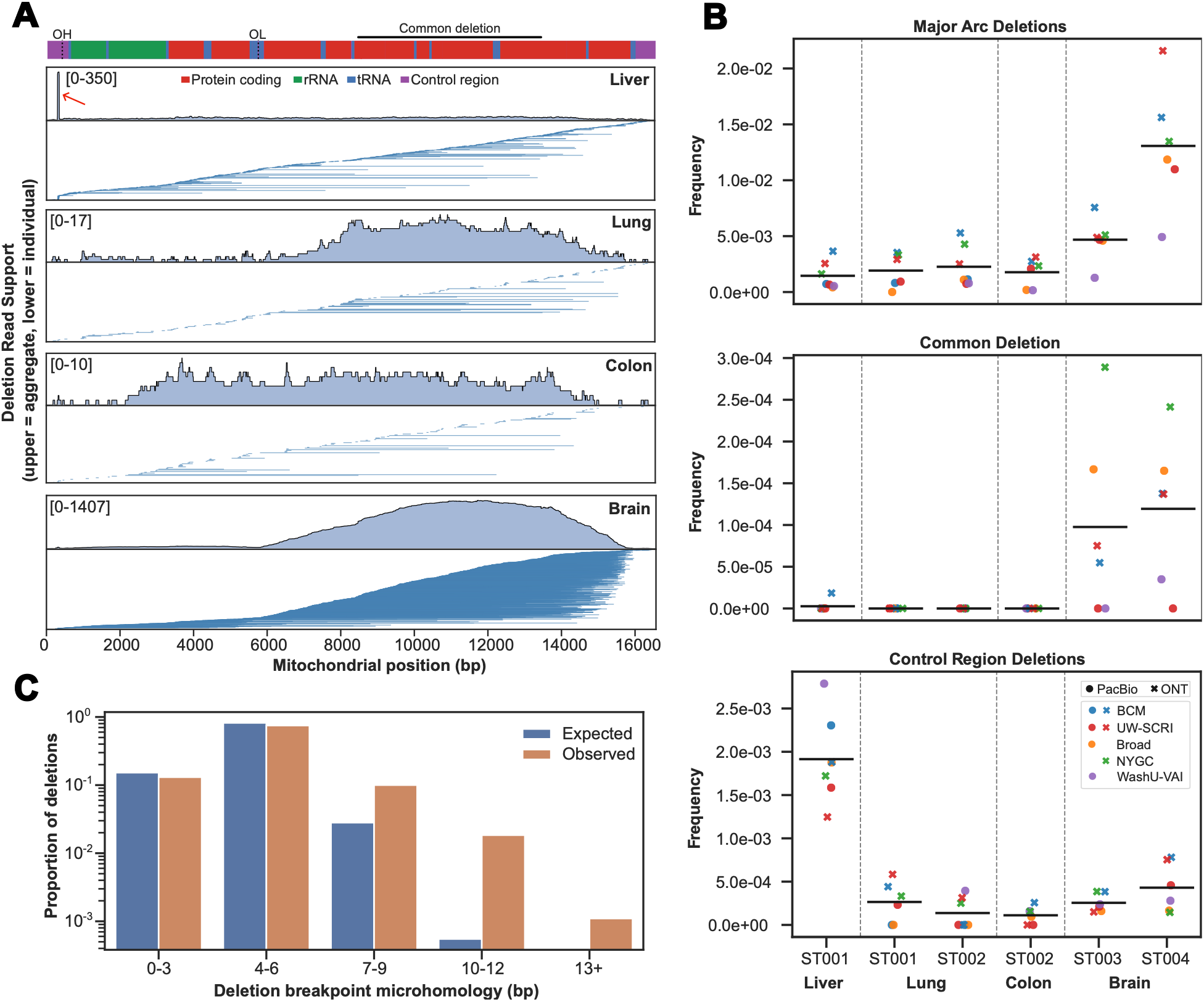
MitoScope uncovers age- and tissue-associated patterns of mitochondrial deletion landscapes. **A** Aggregate and read-level views of mtDNA deletions with associated size and genomic positions. The upper panels show the aggregate deletion depth at each base pair location across the mitochondrial genome with the minimum and maximum deletion-depth indicated in the upper-left corner of each panel. The lower panels show the individual deletion events underlying the aggregate signal. Each horizontal blue line represents one deletion-supporting long read, and its genomic start and end positions indicate the deletion breakpoints and size. Deletions in tissues with multiple donors (Lung and Brain) were aggregated and are shown on a tissue-specific basis. The red arrow indicates a known 50 bp control region deletion in the D-loop (m.298_347del50) identified in ST001 Liver. OH/OL represent the origin of heavy strand and light strand replication, respectively. The canonical 4,977 bp ‘common deletion’ is indicated by the horizontal line above. **B** Frequency of deletions in the major arc region, the ‘common deletion’, and deletions in the control region are shown for donors and tissues across different GCCs and sequencing technologies. Horizontal lines represent mean frequency. **C** Levels of microhomology (bp) between flanking regions surrounding breakpoints in unique deletions across samples (n=3,638) and simulated control deletions (n=18,190, five per observed deletion) are shown with significantly higher levels of observed microhomology compared to expected by chance (P < 0.001, Wilcoxon rank-sum test).

The canonical 4,977 bp “common deletion,” which removes five tRNA genes and seven protein-coding genes and is known to compromise respiratory chain function at high heteroplasmy levels [54], was detected exclusively in brain at low levels (HF ∼0.01%) and was essentially absent in other tissues (**Fig. 4A and B**). Except in liver, 88–97% of deletions mapped to the major arc, whereas a 50 bp control-region deletion in the D-loop (m.298_347del50) predominated in liver (**Fig. 4A and B**). This deletion has previously been reported in the germline of healthy Chinese cohorts [55] and as a recurrent, clonally expanded somatic lesion in gastric adenocarcinoma [56] and aging human putamen [53]; its presence as a major deletion class in liver but absence in matched lung from the same donor suggests a mosaic, tissue-restricted event.

Deletion breakpoints were frequently flanked by short direct repeats or microhomology. The 4,977 bp “common deletion” is flanked by a 13 bp perfect direct repeat and m.298_347del50 by a 9 bp perfect direct repeat. We examined sequence features across all identified deletion junctions (n=3,638) and identified short direct repeats (4–15 bp) across 87% of unique breakpoints. The observed distribution of microhomology was significantly higher than expected by chance (Wilcoxon rank-sum, P < 0.001), where junctions with >6 bp microhomology were ∼4-fold more frequent than expected (**Fig. 4C**). This observed enrichment of short direct repeats and microhomology at deletion junctions identified using MitoScope is consistent with sequence features that have previously been reported at mtDNA deletion junctions [57].

### Long-read detection of NUMTs reveals tissue-mosaic events, underreported and previously unidentified NUMTs, and trinucleotide biases in insertion contexts

NUMTs have been linked to carcinogenesis and to age-related processes involving genomic instability and mitochondrial dysfunction [11], but whether they predominantly act as causal drivers or as byproducts of cellular stress remains unresolved. Additionally, NUMTs’ genomic positions, allele frequencies, and tissue distributions can profoundly affect variant interpretation and nuclear–mitochondrial crosstalk. Building on this, we systematically profiled insertion-derived NUMTs in the nuclear genomes of the benchmarking tissues. Split-read derived NUMTs were excluded from the analysis as they may only have one identified breakpoint, complicating direct comparison with insertion-derived NUMTs. High-confidence NUMTs required support in at least two independent contexts (distinct GCCs, technologies, donors, or tissues), whereas events observed in only a single sample replicate across GCCs and platforms were classified as low-confidence and excluded from downstream analyses. This strategy yielded 63 NUMT calls, of which 57 met high-confidence criteria and collapsed into 17 distinct NUMTs (**Fig. 5A**), indicating a modest number of NUMTs shared across donors and tissues with strong technical support. All 57 high-confidence NUMTs had support from both PacBio and ONT, whereas the six low-confidence calls were technology-specific, with 5 detected only by ONT and one only by PacBio. The high-confidence NUMTs ranged from 32 bp to 870 bp in length (median 66 bp, mean 153 bp, s.d. 203 bp), with insertion breakpoints distributed across several chromosomes (**Fig. 5C**), consistent with the documented landscape of NUMTs with widespread integration across the nuclear genome [11].

**Fig. 5.**
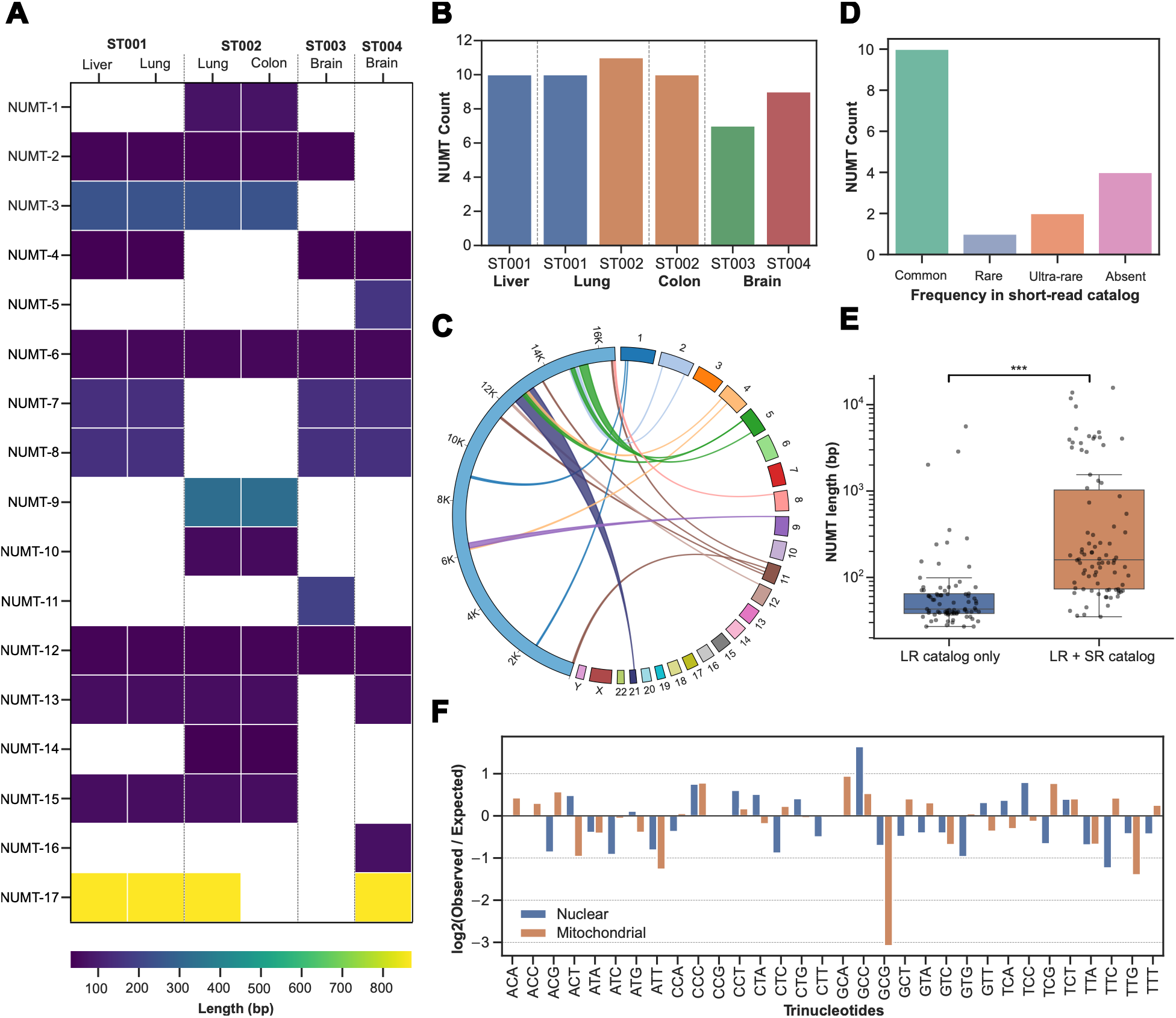
Long-read detection of NUMTs reveals tissue-mosaic events, underreported and previously unidentified NUMTs, and trinucleotide biases in insertion contexts. **A** NUMTs called across benchmarking tissues with color representing the length (bp) of NUMTs. **B** The number of NUMTs identified per tissue across donors. **C** Circos plot showing the nuclear breakpoint of NUMTs with the corresponding aligned section of the mitochondrial genome. Link colors correspond to the nuclear chromosome for the NUMT. **D** The number of common, rare, ultra-rare, and absent NUMTs categorized by frequency groups from the short-read NUMT catalog 100,000 Genomes Project in England. **E** Length distributions of 168 insertion-derived NUMTs identified in the SeqFirst long-read cohort, stratified by representation in the 100,000 Genomes Project short-read catalog. NUMTs absent from the short-read catalog were significantly shorter than shared NUMTs. Data are represented as box- and-whisker plots with the box representing interquartile range (IQR, 25th–75th percentiles), the line within the box representing the median, and whiskers extending to 1.5*IQR (***P < 0.001, Wilcoxon rank-sum test). **F** Trinucleotides surrounding nuclear and mitochondrial breakpoints of NUMTs identified in SeqFirst samples show significantly differing distributions than expected by chance for both the mitochondrial and nuclear genomes (P < 0.001, Chi-square goodness-of-fit).

Across benchmarking tissues, each sample harbored 7 to 11 NUMTs (mean 9.5; s.d. = 1.4) (**Fig. 5B**), an approximately two-fold higher average NUMT count compared to a recent short-read study of 66,083 genomes from the 100,000 Genomes Project [11], where 1,637 unique NUMTs, ranging from 24 bp to the length of the full mitochondrial genome, were identified with an average of 4.7 NUMTs per individual. This difference in average NUMT counts underscores the substantial reservoir of non-reference NUMTs that potentially remain undetected by short-read sequencing and highlights long-read sequencing as a valuable modality for capturing these unique insertion events. For donors with two profiled tissues (ST001 and ST002), all but one NUMT were shared between tissues (**Fig. 5A**), consistent with predominantly germline or early embryonic insertions that are widely distributed across somatic lineages. In contrast, a single NUMT at chr21:9,676,568 (NUMT-17) was detected only in the lung of donor ST002 with variant allele frequencies 0.25-0.32 and absent from the matched colon, indicating a tissue-restricted mosaic NUMT.

Using the population frequencies from the 100,000 Genomes Project short-read study, we classified the 17 high-confidence NUMTs into 10 common (allele frequency ≥1%), one rare (<1% and ≥0.1%), two ultra-rare (<0.1%), and four previously unreported NUMTs that were absent from the 66,083-genome survey (**Fig. 5D**). Surprisingly, all three of the NUMTs categorized as rare or ultra-rare according to the short-read study were present in at least two or more of the four benchmarking donors, implying that many of these ‘rare/ultra-rare’ NUMTs may be considerably more common than previously reported in short-read based surveys.

To expand NUMT discovery beyond the benchmarking tissues and incorporate a more comprehensive long-read NUMT catalog, we leveraged long-read whole-genome data from 359 individuals enrolled in the SeqFirst-neo project [58]. Across the long-read cohort, we identified 168 unique non-reference NUMTs, collapsed across calls detected by both PacBio and ONT (87%) and technology-specific calls (4% ONT only, 9% PacBio only). To investigate properties of NUMTs differentially represented in long- and short-read datasets, we compared these 168 NUMTs identified in the long-read cohort with the 100,000 Genomes Project short-read catalog. Overall, 82 (49%) of the 168 unique NUMTs were absent from the existing short-read catalog (**Additional file 2: Table S10)**. Notably, 59 of the 82 long-read-specific events were rare (allele frequency <1%) among the SeqFirst cohort. Additionally, NUMTs absent from the short-read catalog were significantly shorter than NUMTs represented in both catalogs (Wilcoxon rank-sum, P < 0.001), with 88% (n=72) of long-read-specific NUMTs measuring less than 100 bp (**Fig. 5E**). The preferential identification of short, low-frequency NUMTs by long-read sequencing may reflect challenges of resolving short mitochondrial-derived insertions from short-read data, particularly when sequence fragments cannot be uniquely anchored to nuclear flanking DNA or distinguished from mitochondrial sequence. As the long-read and short-read catalogs were generated from different cohorts and analytical strategies, these results reflect differences in catalog representation rather than a direct comparison of NUMT burden or platform sensitivity. The identification of NUMTs absent from existing short-read catalogs, particularly shorter and low-frequency NUMTs, highlights the need for more comprehensive, publicly accessible long-read-based NUMT catalogs to improve variant interpretation, mitigate NUMT-mediated artifacts, and enable more accurate studies of nuclear–mitochondrial genome interactions.

To examine sequence features surrounding NUMT insertion sites, we analyzed the trinucleotide context surrounding NUMT breakpoints in both the nuclear and mitochondrial loci of the SeqFirst cohort (**Fig. 5F**). The observed trinucleotide distribution at NUMT breakpoints deviated markedly from the expected genome-wide frequencies in both the nuclear and mitochondrial compartments (P < 0.001). Specifically, breakpoints were enriched for nCC/CCn motifs and depleted for nTT/TTn motifs across both nuclear and mitochondrial loci. In nuclear breakpoints, nCG trinucleotides were also underrepresented, while mitochondrial breakpoints exhibited the strongest depletion for GCG (>3-fold reduction relative to background). These results indicate that sequences surrounding nuclear and mitochondrial sites associated with NUMT integration exhibit nonrandom local sequence composition, consistent with previous reports of sequence biases at sites of mtDNA integration [11].

## Discussion

In this work, we present MitoScope, a scalable long-read workflow for high-fidelity detection and characterization of mtDNA variation, including low-frequency heteroplasmies, large deletions, and NUMTs, across SMaHT benchmarking tissues and expanded cohorts. MitoScope demonstrates high sensitivity and precision for mtDNA variant detection in HapMap mixture cell lines and enables characterization of mtDNA variation at both bulk and read-level resolution. Across human tissues, MitoScope successfully identifies established patterns of somatic mtDNA variation, including tissue- and donor-specific heteroplasmic variants, age-associated accumulation of mtDNA deletions, strand- and sequence-context biases, and low-frequency disease-associated heteroplasmies in clinically unaffected tissues. MitoScope identifies NUMTs in long-read data that were absent from existing short-read catalogs and enriched for shorter and low-frequency NUMTs. These findings demonstrate the ability of MitoScope to uncover and characterize established and previously underrepresented patterns of mitochondrial and nuclear-mitochondrial sequence variation, providing a scalable framework for future studies of mtDNA variation across aging and disease.

The tissue analyses provide a biologically relevant benchmarking resource that extends validation beyond the blood- and cell-line-based datasets that have dominated many prior mtDNA studies. By analyzing postmortem donor tissues with matched PacBio, ONT, high-depth Illumina WGS, and duplex-sequencing data generated across multiple centers, we demonstrate the reproducibility, limitations, and complementary value of MitoScope across orthogonal technologies while showing that it recovers established features of mtDNA variation in human tissues. While the cohort size, bulk-tissue design, limited tissue sampling, and absence of lineage-resolved data preclude inference about the causes or consequences of the observed biological patterns, this multi-platform tissue resource provides a foundation for future, larger, lineage-informed, deeply phenotyped, and longitudinal studies designed to test the mechanisms and clinical consequences underlying tissue-specific and age-associated mtDNA heteroplasmy, deletions, and NUMT variation.

We demonstrate that MitoScope can detect and report large deletions at read-level resolution and can identify individual mtDNA molecules containing both SNVs and deletions. Given the low heteroplasmy of the evaluated SNVs and the limited number of reads with co-occurring events, we did not attempt to construct clonal relationships between SNVs and deletions. However, we do establish a workflow for future analyses of deletion-bearing mitochondrial haplotypes, molecular co-occurrence patterns, and clonal mtDNA evolution in larger cohorts with deeper mtDNA coverage. Mutations in clonal hematopoiesis genes recently shown to have associations with high mtDNA burden (TERT, TCL1A, and SMC4) [9] should also be further investigated for somatic driver mutations. Initiatives like the SMaHT Network, with its standardized multi-center long-read benchmarking across diverse post-mortem tissues, enable unprecedented systematic surveys of mtDNA mosaicism using robust tools like MitoScope and optimized protocols. Such expanded studies will facilitate critical phenotype-genotype associations to elucidate these relationships. These efforts are expected to illuminate variation patterns; uncover hidden heteroplasmies, deletions, and NUMTs; and reveal their contributions to tissue-specific instabilities, late-onset disorders, variant evolution, and nuclear-mitochondrial crosstalk.

A major challenge in long-read mtDNA analysis is the marked variability in inferred copy number, which significantly underestimates true abundance compared to short-read estimates due to protocol-specific biases in high molecular weight DNA preparation. Size-selected libraries favoring long fragments preferentially lose short mtDNA molecules during shearing and selection, as these circular ∼16.5 Kbp genomes fragment inefficiently and are discarded, systematically biasing all long-read datasets analyzed here. This underrepresentation not only depresses apparent mtDNA coverage but also risks uncovering low-frequency somatic variants and large mtDNA deletions, as reduced input read depth amplifies stochastic sampling effects and diminishes statistical power for rare alleles. Standardized, mtDNA-inclusive library prep protocols or separate enrichment strategies will be essential to mitigate these artifacts and ensure accurate quantification of mtDNA variation in future long-read studies.

We also recognize that mtDNA methylation analysis represents an important and exciting frontier for understanding epigenetic regulation of mitochondrial function and nuclear-mitochondrial interactions. While MitoScope can generate single-read 5mC methylation calls across mtDNA CpG sites from long-read data and enable nucleotide-resolution epigenetic profiling alongside variant detection, we did not extensively analyze mtDNA methylation due to its controversial nature and pronounced technical discordance between platforms—PacBio calls exhibited higher noise and elevated 5mC levels inconsistent with reports of very low mitochondrial methylation [59], while ONT data aligned better with expected near-background patterns. This variability likely reflects platform-specific base calling artifacts or limited CpG contexts in the compact mtDNA, underscoring the need for orthogonal validation in future studies to resolve whether trace methylation plays a functional role in mtDNA regulation.

## Conclusions

MitoScope is a containerized end-to-end long-read workflow enabling high-fidelity detection of somatic mtDNA variants, structural deletions, and NUMTs across human tissues. Benchmarking demonstrates high sensitivity and precision with heteroplasmy estimates concordant with bulk short read and duplex sequencing data. Applied to the SMaHT Network benchmarking tissues, MitoScope successfully uncovers previously established biological patterns of mtDNA, including low-frequency pathogenic heteroplasmies in clinically normal tissues, age- and tissue-associated accumulation of mtDNA mutations including large deletions, and enriched deletion breakpoints near regions of microhomology. NUMTs derived from long-reads reveal events that are absent from existing short-read catalogs, exposing a potential hidden reservoir of nuclear-mitochondrial variation and providing a framework for future studies across matched short-read and long-read cohorts. Expanding MitoScope analyses to larger, deeply phenotyped cohorts will be essential to disentangle germline from somatic variation and to establish the phenotypic consequences of subclinical mitochondrial mosaicism.

## Methods

### MitoScope mtDNA selection

BAM/CRAM files are converted to FASTQs prior to k-mer selection. A k-mer count database of mtDNA k-mer sequences (k= 29) is generated using the human mitochondrial reference sequence (GenBank: NC_012920.1) and the k-mer tool kit, KMC3 (v3.2.1) [60]. The k-mer count database is used to selectively retain reads in the FASTQ file with >2,500 matching k-mers for retention of reads with sufficient length and sequence similarity to mtDNA. After k-mer selection, reads are aligned to the mitochondrial genome using Minimap2 (v2.24) [61] and evaluated for the removal of contaminating NUMT reads. Thresholds are enforced for maximum soft-clipped sequence in a read that does not align back to the mitochondrial genome as well as the fraction of 5mC methylated CpG sites in a read. Up to 200 bp of unaligned soft-clipping and a maximum of 50% methylated CpG sites are used for the removal of NUMTs to avoid compromising mtDNA read counts. A stricter threshold of 200 bp of unaligned sequence alongside a more lenient threshold of 50% methylated CpG sites is used to prioritize removal of reads with significant levels of unaligned sequence while also discarding reads with good alignment but deviating methylation profiles. A methylation likelihood probability threshold of 0.5 is used to classify sites as methylated or unmethylated. Reads failing either filtration threshold are filtered out and written to a separate discard BAM file. Foldback artifacts, where a molecule folds back on itself and appears as a split read aligning to opposite strands, are also removed to avoid compromising assembly accuracy.

### MitoScope variant calling

MitoScope uses baldur [29] (git-commit: 73cb26e) to call heteroplasmic SNVs, indels, and large deletions (options: -P 5 -q 20 -Q 20 -I 30 --output-deletions --small-deletion-limit 25 --large-deletion-limit 25 --indel-thresholds 0.1 0.1 --snv-thresholds 0.0005 0.0025). Structural variants are called using Sniffles2 [30], (v2.6.2) with parameters customized to identify heteroplasmic SVs (options: --qc-output-all --minsvlen 5 --minsupport 4). Variant Effect Predictor (VEP) [31] (v115.0) is run on the resulting SNV/indel VCF for variant annotations and MITOMAP [32] annotations are integrated using BCFtools (v1.21) [62] for pathogenicity and population frequencies. Read by variant matrices are generated recording the presence or absence of SNVs and indels across individual reads. Haplotypes are encoded as ‘0’ for the reference allele, ‘1’ for the target alternate allele, ‘2’ for another alternate allele, and ‘-1’ for no coverage at site. A separate file is generated recording large deletions (>45) with their genomic coordinates across individual reads.

### MitoScope assembly generation

Mitochondrial genome assemblies are generated using meta-flye [35] (v2.9.6, options: --meta -m 2500). Meta-flye is run using an initial set of random downsampled reads (n=100) generated by seqtk (v1.5). If the expected ∼16.5 Kbp circular contig is not obtained, another random subset of reads is used to rerun assembly. This is repeated until a successful assembly is generated or a maximum number of iterations is reached (default=10). A tolerance of 100 bp relative to the length of the mitochondrial genome (16,569 bp) is allowed for an assembly to be considered the valid length. For low-coverage samples where a circular assembly of the expected length cannot be achieved, the retained assembly output is from the final iteration.

### MitoScope copy number calculation

mtDNA copy number (CN) is estimated using coverage of the mitochondrial genome and the nuclear genome (chr1-22,X,Y). Coverage values are generated using mosdepth [36] and the relative mtDNA CN per cell, assuming a diploid nuclear genome, is calculated as (mtDNA coverage / nuclear coverage * 2).

### MitoScope NUMT characterization

Non-reference NUMTs are called based on insertion calls or split read-derived calls. For insertion-derived NUMTs, structural variants are called using Sniffles2 [30] (v2.6.2, options: --minsvlen 20) on input CRAM files aligned to the GRCh38 reference genome. Only insertions with at least two supporting reads are retained for consideration as NUMT candidates. Sequences for called insertions are written to a FASTA file that is mapped against the mitochondrial reference genome (revised Cambridge Reference Sequence) using BLAST [63] (v2.16.0, options: -word_size 25 - perc_identity 95). Insertions that share >95% sequence identity with mtDNA are run through BLAST a second time using the GRCh38 reference to retain only those insertions with higher sequence similarity to the mitochondrial genome than to the nuclear genome. Split read-derived NUMTs are identified from the same aligned CRAMs (to GRCh38) based on reads with supplementary alignments to both the mitochondrial (chrM) and nuclear genomes (chr1-22,X,Y). Only split-read derived NUMT candidates with breakpoints with minimum support from at least 4 different reads are retained and reported. Final reported NUMTs are categorized according to their nuclear and mitochondrial breakpoint locations, levels of read support, and variant allele frequencies.

### SMaHT benchmarking data

The sequencing data for the HapMap mixture cell line and the benchmarking tissues (ST001-ST004) was generated by the Somatic Mosaicism across Human Tissues (SMaHT) benchmarking effort [20–22]. Long-read (PacBio and ONT) and short-read (Illumina) DNA sequencing data was downloaded from the SMaHT data portal in the form of aligned and indexed CRAM files generated using the GRCh38 reference. The HapMap mixture sample and all tissue samples were each sequenced by at least three different GCCs (BCM, UW-SCRI, WashU-VAI, Broad, NYGC). The downsampled HapMap mixture samples with 30x, 60x, and 90x coverage were generated by the BCM GCC separately for ONT and PacBio data. Duplex sequencing data for the benchmarking tissues was generated by the Broad GCC using the CODEC duplex assay. Intact tissue or homogenates were used as starting input material. Further details on sequencing and library preparation of the HapMap and benchmarking samples can be found in the SMaHT data portal, the SMaHT HapMap mixtures paper, the SMaHT duplex benchmarking paper, and the SMaHT flagship paper [21,22,47].

### Benchmarking tool performance using the HapMap mixture cell line data

The mitochondrial truth set used to evaluate variant calling performance was derived from the HPRC Version 2 release of the pangenome-graph derived variant calls from the minigraph-cactus pipeline (hprc-v2.0-mc-grch38.wave.vcf.gz). The truth set was generated by subsetting the VCF to variants located on chrM and present in at least one of the HapMap mixture samples (HG005, HG02622, HG02486, HG02257, HG002, HG00438). As the variant calls made on the HapMap mixture samples were indistinguishable based on sample origin, we generated a collapsed set of HapMap variant calls (98 SNVs, 13 indels) from the truth set. Genotype columns in the mitochondrial truth set VCF corresponding to each HapMap sample in the mixture were collapsed to a single column indicating variant presence. Variant calling performance was evaluated across Mitoscope [25] (v0.4.0), Himito [18] (v1.1.2), and Mitorsaw [19] (v0.2.14) initially using default parameters and separately using matched minimum VAF settings (--vaf_threshold 0.0025 for Himito and --minimum_maf 0.0025 for Mitorsaw). Similar to MitoScope’s separate minimum VAF threshold for indels set to a higher value of 0.1, Himito also enforces a separate VAF threshold for indels (--indel_false_threshold) that also defaults to 0.1. GATK Mutect2 [64] (v4.5.0.0) was run in mitochondrial mode (--mitochondria-mode) with otherwise default parameters followed by FilterMutectCalls to retain passing variants. Sensitivity, precision, and F1 scores were calculated using RTG tools (v3.12.1) [65].

### Variant calling of CODEC duplex sequencing data

CODEC duplex sequencing data downloaded from the SMaHT data portal was processed using CODECsuite [66] (v1.1.5) and variant calling was performed using Minipileup [67] (v1.4b). The CODECsuite filter function was run using the following custom parameter settings: -m 30 -q 20 - Q 0.7 -N 0.05. Minipileup was run using the following custom parameter settings: -vcC -q30 -Q20 -s 4 -a 2 -T 2.

### SNV/indel analysis on benchmarking tissues

MitoScope SNV and indel variant calls were generated using PacBio data (merged across GCCs) and ONT data (merged across GCCs). Variant calls between platforms were collapsed and heteroplasmy fractions were averaged to generate a single set of MitoScope variant calls. The status of PacBio and ONT support for each variant call was retained for further comparisons with the bulk short-read and duplex variant calls. MitoScope variants calls that had support across both PacBio and ONT data or support from the duplex or bulk short-read data were retained as the high-confidence set for further downstream analysis. MitoScope variant calls that were detected only in PacBio or only in ONT with no additional support in duplex or bulk short-read data were considered lower confidence. Heavy- and light-strand base substitutions were assigned according to the reference allele in the revised Cambridge Reference Sequence (rCRS), which is reported in the light-strand orientation. SNVs with pyrimidine reference alleles (C or T) were classified as light-strand substitutions. For SNVs with purine reference alleles (A or G), the complementary pyrimidine substitution was classified as a heavy-strand substitution. Binomial tests were performed to assess whether observed variant counts were significantly enriched or depleted across mitochondrial gene regions (control region (hypervariable and outside hypervariable regions), protein coding, rRNA, tRNA) using expected counts proportional to the length of each region. Variant counts were aggregated per region for testing, and p-values were adjusted for multiple testing correction using the Benjamini–Hochberg false discovery rate method. For visualization, variant counts were normalized by region length to obtain variant frequencies.

### Deletion analysis

Frequency and breakpoint spanning deletion analysis was performed using output from baldur with Sniffles2 calls used for further validation of the m.298_347del50 control region deletion and the 4,977 bp ‘common’ deletion. Only deletions >45 bp were kept for quantification. Major arc deletions were quantified by counting deletions with breakpoints falling within m.5576-15976. The “common deletion” was quantified by counting deletions whose start/end breakpoints fell within m.8460-8480 and m.13430-13455, respectively. Control region deletions were quantified by counting deletions with breakpoints within m.16024-16569 or m.1-576. The m.298_347del50 control region deletion was quantified for deletions with start/end breakpoints within m.287-307 and m.336-356. Frequencies for each deletion group were calculated by normalizing read counts with deletions by total mtDNA read counts. Deletion breakpoint microhomology analysis was performed by identifying perfect direct repeats between the first 20 bp of deletion sequence and the 20 bp of flanking sequence at the end of the deletion using BLAST (options: -word_size 4 - perc_identity 100). For each reported deletion with unique breakpoints across the benchmarking tissues, control data was simulated with matching deletion length and randomized breakpoints with five simulated deletions per observed deletion (3,638 observed deletions, 18,190 simulated deletions). Wilcoxon-rank sum test was used to test for statistical significance between observed and expected distributions.

### NUMT analysis

NUMTs included in the analysis of the benchmarking tissues and SeqFirst samples were insertion-derived. Split read-derived NUMTs reported by the workflow were not included to avoid the complication of comparing NUMTs with potentially only one mitochondrial breakpoint. Precise nuclear and mitochondrial NUMT coordinates for benchmarking tissue samples with the potential to identify individuals were not specified for participant privacy. For comparison of NUMTs across different samples, a tolerance of 100 bp was used to cluster NUMTs by nuclear and mitochondrial breakpoints. SeqFirst samples with both PacBio and ONT data (n=359) were used to derive the long-read NUMT catalog. NUMTs that were called across either or both technologies were included in the set of NUMTs and classified the same way as in the benchmarking tissues using nuclear and mitochondrial breakpoints. The short-read NUMT catalog consisted of 1,637 NUMTs from the 100,000 Genomes Project in England [11]. NUMT comparisons between the different catalogs were performed using 100 bp tolerance surrounding nuclear breakpoints. Trinucleotide breakpoint analysis was performed using the directly flanking trinucleotide sequences surrounding the nuclear and mitochondrial breakpoints of NUMTs. Breakpoints directly at the beginning or end (1 or 16,569) that could pose issues with genome wrap-around were excluded. Genome-wide trinucleotide distributions were used for the control set by calculating the proportion of each trinucleotide across the mitochondrial genome and the nuclear genome (chr1-22,X,Y). Trinucleotides sequences were collapsed and counted together with their corresponding reverse complements. Chi-square goodness of fit test was used to test for significance between observed and expected trinucleotide distributions.

## DECLARATIONS

### Ethics approval and consent to participate

Ethical approval for the tissue collection was obtained by National Disease Research Interchange (NDRI) through the University of Pennsylvania (IRB#5 FWA00004028). Frozen dorsolateral prefrontal cortex tissue from deceased donors ST003 and ST004 was obtained from the NIH NeuroBioBank repository at the University of Maryland Brain and Tissue Bank (UMBTB). Collection of brain tissue from donors was performed as approved by the IRB of the University of Maryland, Baltimore and the University of Miami School of Medicine where the respective cases were recovered.

### Consent for publication

Not applicable.

### Availability of data and materials

This study was conducted as part of the NIH Common Fund consortium initiative, Somatic Mosaicism across Human Tissues (SMaHT). The benchmark datasets described in this study are available through dbGaP under the study accession number phs004193 and from the SMaHT data portal [68]. The data used in this work was provided by the SMaHT Data Analysis Center (DAC) [1UM1DA058230] on behalf of the SMaHT Network. More information about the SMaHT Network is available online [69]. The HPRC pangenome-graph derived variant calls used for the mitochondrial truth set are available from the HPRC data portal [70]. The PacBio and ONT SeqFirst-neo data used for NUMT analysis are available through the GREGoR Consortium: Genomics Research to Elucidate the Genetics of Rare Disease dbGaP study (accession no. phs003047) in NHGRI’s Analysis Visualization and Informatics Lab-space (AnVIL) platform [71]. MitoScope code is publicly available on GitHub [25] under an MIT license and the version of MitoScope used in this manuscript is available on Zenodo [72] under the same license. Analysis scripts used to create and summarize results have been deposited in GitHub [73] and are publicly available as of the date of publication. Any additional information required to reanalyze the data reported in this work is available from the lead contact upon request.

### Author contributions

C.Z., J.D.S., C.H.W., and C.L.W. conceptualized the project. C.Z. and C.H.W. performed tool development. C.Z. conducted bioinformatics analyses. C.Z., J.D.S., and C.L.W. wrote the manuscript with input from all authors. C.D.F., E.R., S.R.M., M.R., J.M.W., K.M.M., K.H., T.M., Y.K., J.O., S.N., M.S., A.M., J.T.B., A.B.S., and E.E.E. reviewed and/or edited the manuscript. C.L.W. supervised the project. All authors reviewed and approved the final manuscript.

### Corresponding author

Further information and requests for resources and tools should be directed to and will be fulfilled by the corresponding author, Chia-Lin Wei.

## SUPPLEMENTARY INFORMATION

**Additional file 1:** Supplementary figures S1 to S7 **Additional file 2:** Supplementary tables S1 to S10 **Supplementary Tables:**

**Table S1.** List of software and containers used in MitoScope workflow

**Table S2.** List of MitoScope expected output files

**Table S3.** HapMap QC metrics for MitoScope PacBio and ONT

**Table S4.** HapMap QC metrics for Illumina

**Table S5.** Variant calling performance of MitoScope, Himito, Mitorsaw, and Mutect2 on HapMap samples with default minimum VAF settings

**Table S6.** Variant calling performance of MitoScope, Himito, Mitorsaw, and Mutect2 on HapMap samples with matched minimum VAF settings

**Table S7.** Concordance of heteroplasmy levels between MitoScope and Mutect2 calls for HapMap samples

**Table S8.** Benchmarking tissue QC metrics

**Table S9.** High-confidence heteroplasmic SNVs in benchmarking tissues

**Table S10.** Catalog of unique NUMTs identified in SeqFirst samples

