## Supplemental Figures for "MitoScope: a comprehensive mitochondrial genome analysis workflow for long-read sequencing data"

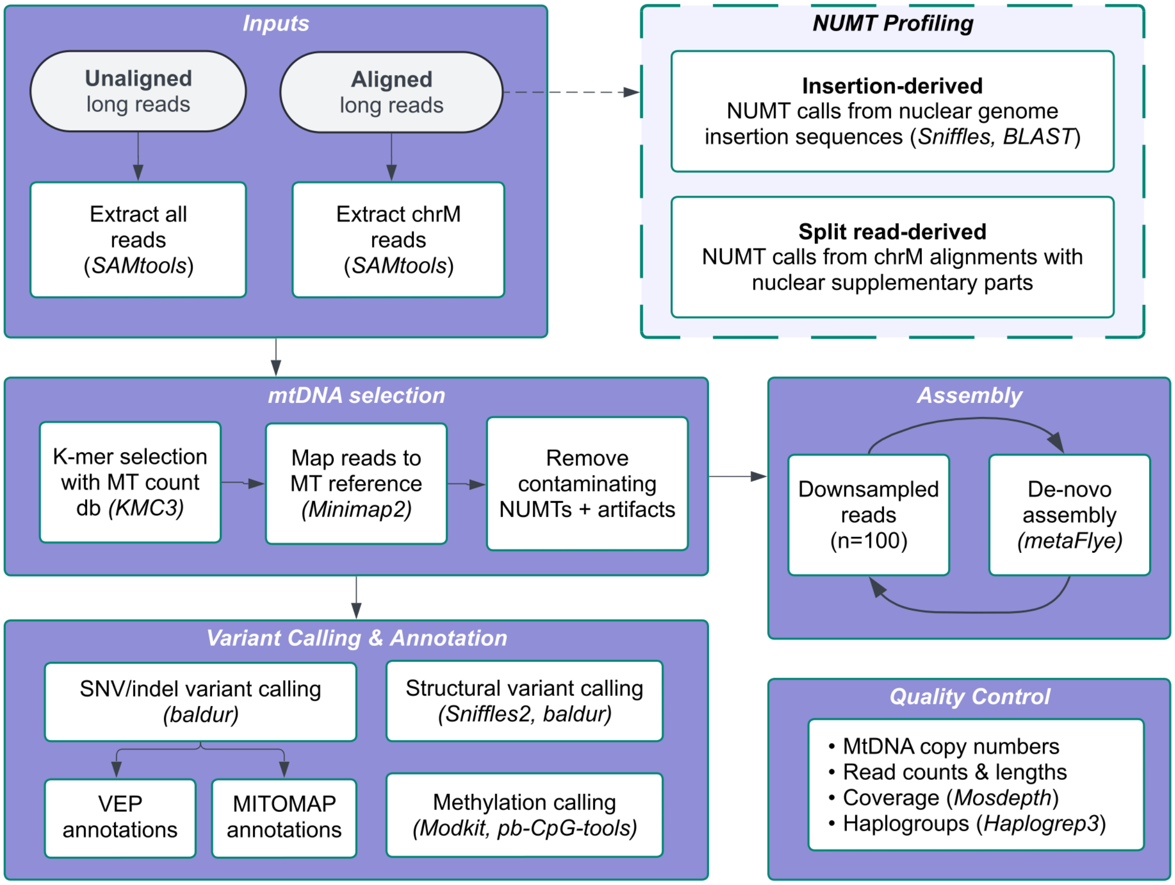


**Fig. S1 MitoScope: A scalable long-read workflow for mtDNA variant detection and NUMT profiling.** MitoScope is built as a fully containerized Nextflow workflow using Singularity images. The core modules that constitute the workflow are: **1)** mtDNA selection, **2)** variant calling and annotation, **3)** assembly generation, **4)** quality control, and **5)** optional NUMT profiling. Sequencing data is accepted as raw unaligned or genome aligned input in the form of FASTQ, BAM, or CRAM files. Relevant open-source tools integrated into the workflow are designated in parenthesis.

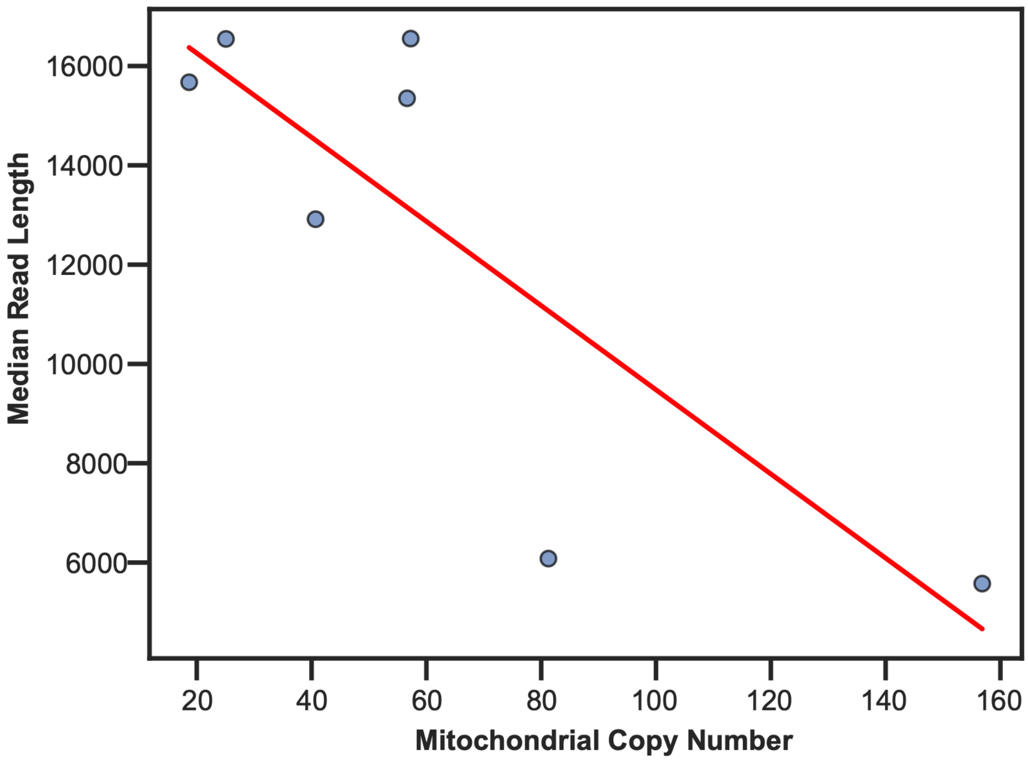


**Fig. S2 HapMap mixture samples reveal an inverse correlation between read lengths and mitochondrial copy number. A** Long-read data for the SMaHT HapMap mixture samples was processed using MitoScope and an inverse correlation between mitochondrial copy number and median read length of mtDNA reads was observed associated with variation in library preparation protocols across different GCCs.

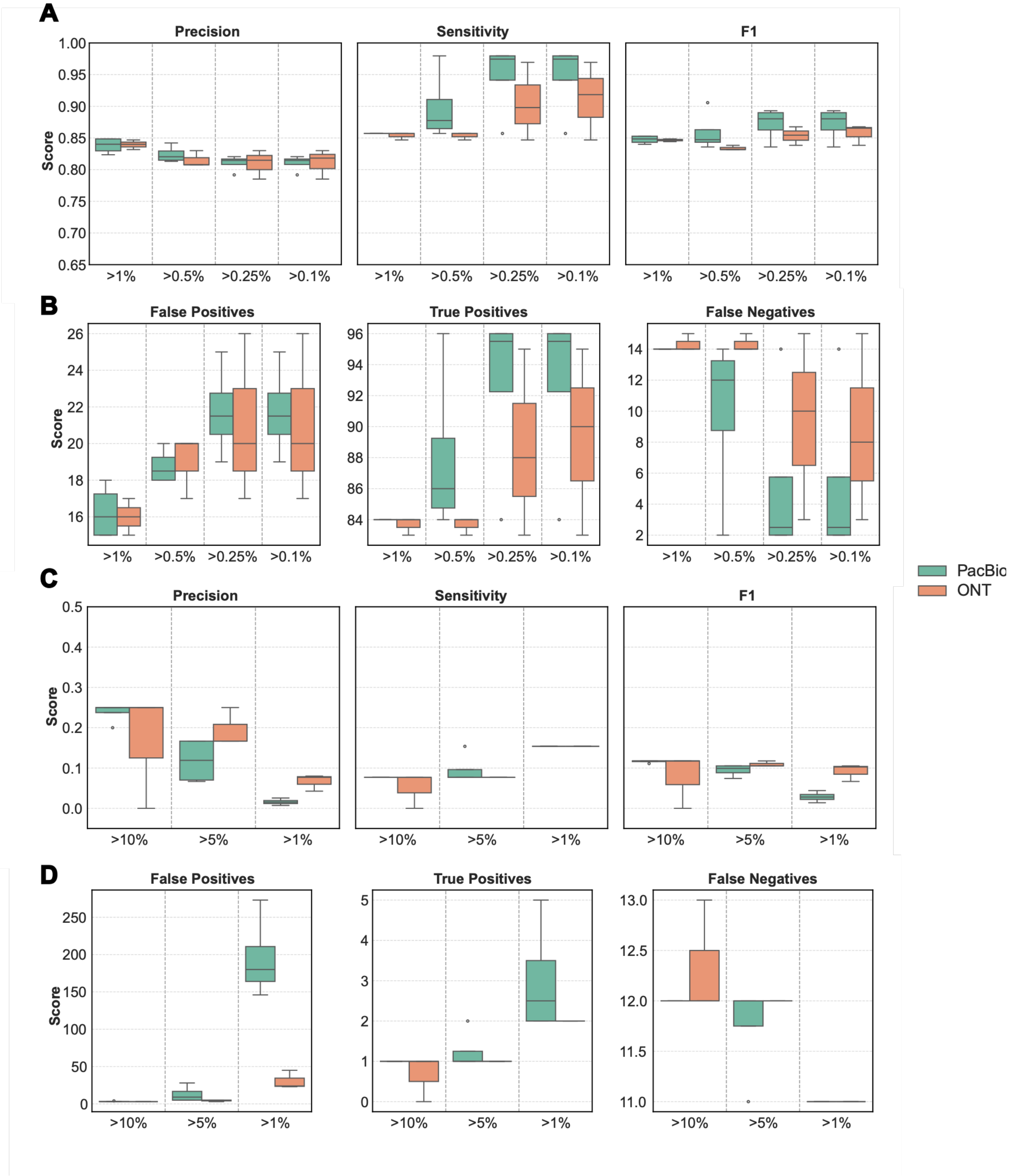


**Fig. S3 Comparison of MitoScope variant calling performance across different minimum VAF thresholds.** Precision, sensitivity, and F1 scores were calculated for SNV (**A**) and indel (**C**) variant calling results from MitoScope using different minimum VAF thresholds. The corresponding counts for false positive, true positive, and false negative calls are also shown for SNVs (**B**) and indels (**D**). Data are represented as box-and-whisker with the box representing interquartile range (IQR, 25th–75th percentiles), the line within the box representing the median, whiskers extending to 1.5*IQR, and outliers depicted using individual points (n=3 GCC replicates for ONT and n=4 for PacBio).

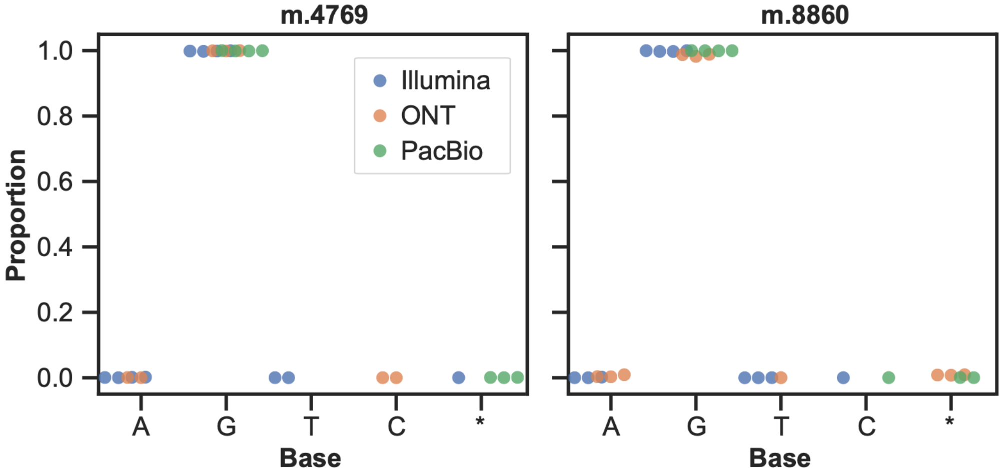


**Fig. S4 HapMap pileup results confirm discordant variant calls from Mutect2.** Pileup base call results for m.4769 and m.8860 where Mutect2 falsely identified heteroplasmy frequencies of 0.7-0.8 while MitoScope correctly identified both variants as homoplasmic with support from the pileup results across Illumina, PacBio, and ONT reads. Deletion of reference base is denoted by an asterisk (*).

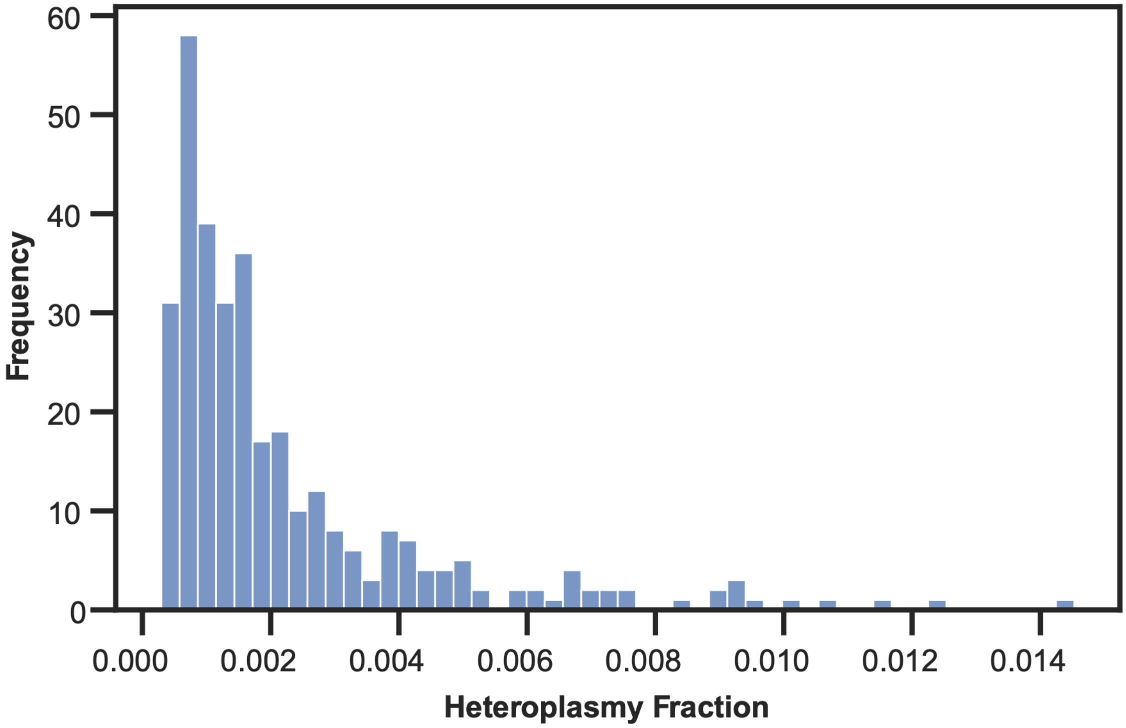


**Fig. S5 Distribution of heteroplasmy levels for variants that were unique to duplex sequencing.** Heteroplasmic SNVs that were only represented in the duplex sequencing data (n=312) were more likely to be ultra low-frequency calls.

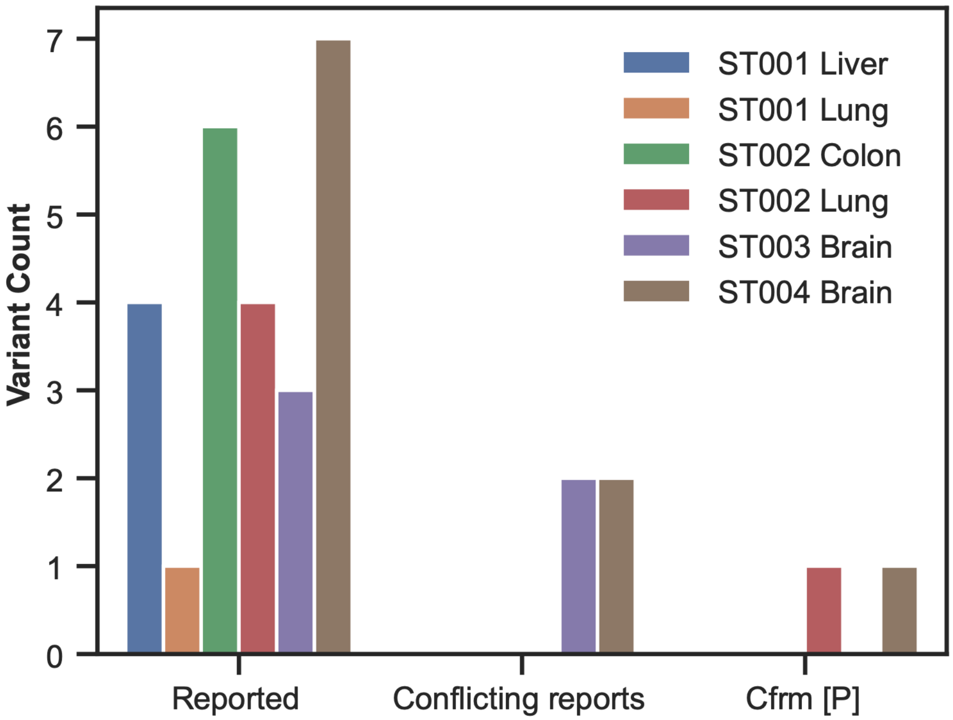


**Fig. S6 SNVs in MITOMAP with reported or confirmed disease associations.** MITOMAP classifies variants as “reported,” with “conflicting reports,” or as “confirmed” (Cfrm) disease variants, with the latter further annotated according to the ClinGen Pathogenicity Ratings (VUS - Variant of Uncertain Significance; LP - Likely Pathogenic; P - Pathogenic).

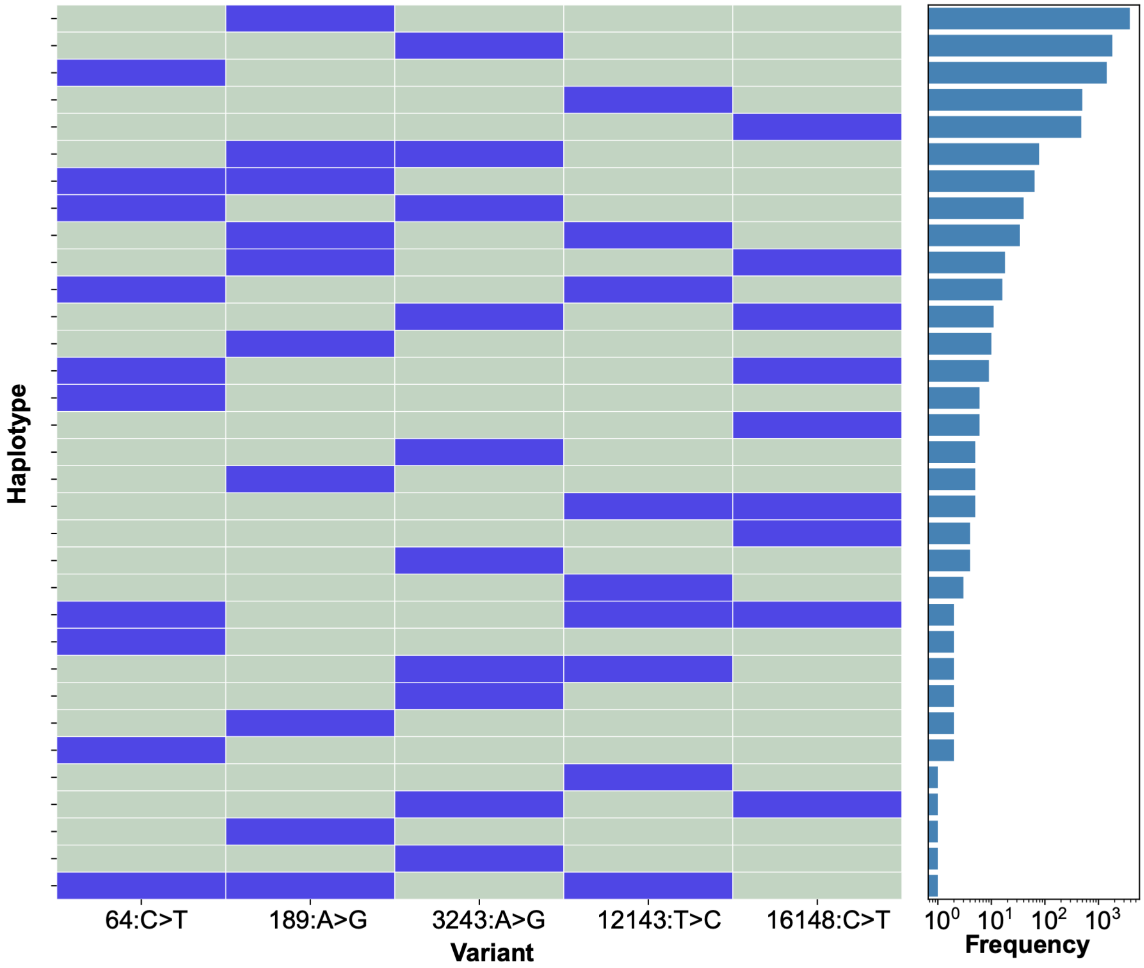


**Fig. S7 Single-molecule haplotypes of heteroplasmic SNVs in ST004 Brain.** Heatmap of distinct haplotypes across five heteroplasmic SNVs detected by both PacBio and ONT. Reads spanning all five sites were combined across platforms. Rows indicate read-level haplotypes; columns indicate variant positions. Dark blue, alternate allele; light green, reference allele. The bar plot indicates read counts per haplotype. Most variant-containing reads carried one heteroplasmic SNV, whereas a minority carried two or three co-occurring variants.
